## Supplementary Information (notes 1-4, figures and tables) for "Differential richness inference for 16S rRNA marker gene surveys" (from another version of this preprint: version 1, 2021-11-08)

### Supplementary Note 1

### 1 Supplementary Note 1 : The type I error rate of sequence similarity thresholds.

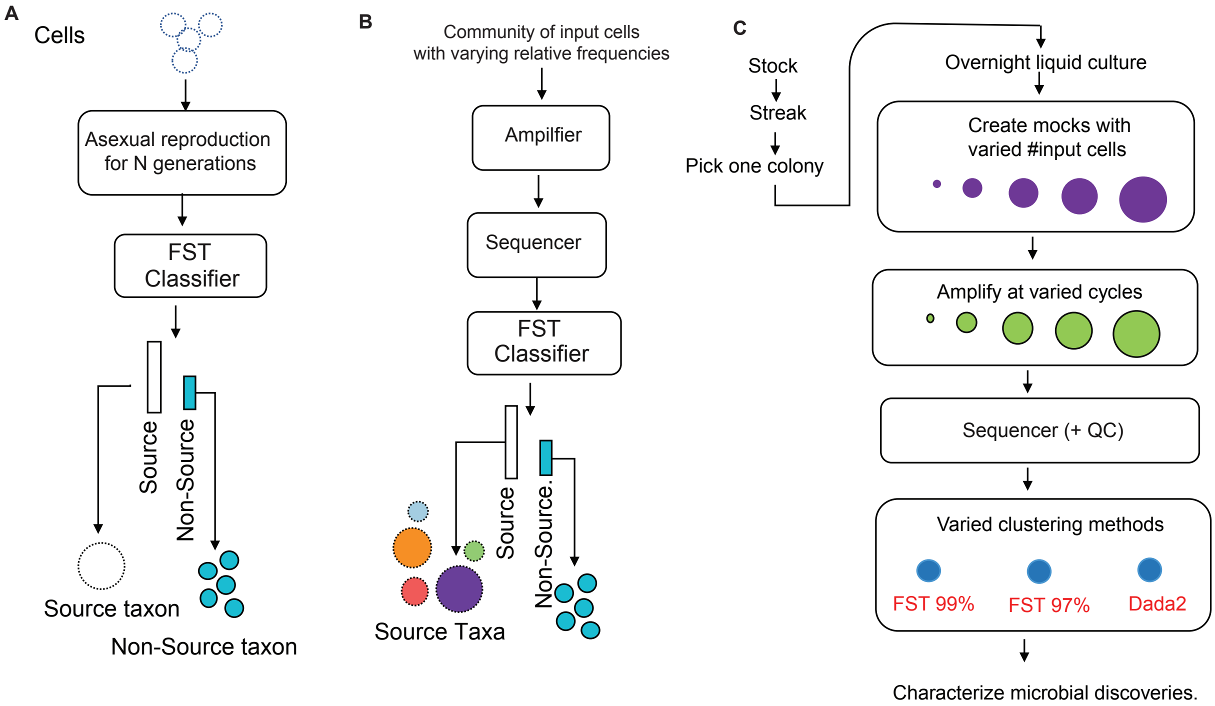


**Figure 1: Questions asked in sections 1.1 - 1.2 and main text section 2.1.** When an apriori fixed sequence similarity threshold (FST) based classifier is input a source (reference) sequence, and a set of its noisy realizations from (A) cellular reproduction-like processes, the type I misclassification errors transform to false microbial discoveries through sequence clustering. We quantify the type I classification error rate. In (B), we seek the same for a community of sequences input to a chain of amplification and sequencing processes. In (C), we aim to experimentally study false discovery accumulations in a 16S survey of single colony derived bacterial culture (main text). The laboratory samples are constructed by varying two experimental parameters (number of input cells and 16S rRNA amplification cycles), and a bioinformatics parameter, the microbial population reconstruction approach.

Given a query and a reference sequence of the same length $L$, we quantify the sequence similarity between the two as the number of sites where they differ in nucleotide identitities. We use the term $FST(\delta)$ to denote a binary sequence classifier that classifies a query as a source type (i.e., reference) when the sequence similarity between the query and source is $\leq \delta L$. Thus, $\delta = .01$ and $\delta = .03$ correspond to 99% and 97% sequence similarity thresholds respectively.

**Fig. 1** outlines a subset of questions asked in this paper. Briefly, given a reference DNA segment and a well-defined set of its noisy realizations as input, we seek the type I error rate of a fixed sequence similarity threshold based classifier $FST(\delta)$. The set of erroneous variants are generated thorugh nucleotide substitutions from (A) asexual cellular reproduction, (B) polymerase chain reaction induced amplification, (C) sequencing machine and (D) a chain of amplification and sequencing processes. In each case, the cardinality of the generated variant set dictates the total number of classifications, and hence the number of type I errors generated. The type I errors generated through these mechanisms go onto organize as false microbial discoveries in a study via the clustering algorithm used.

##### 1.1 Type I error rate of an FST classifier when genotyping natural variation induced by cell growth

Consider a selectively neutral DNA segment of length $L$ in a cell undergoing perfect cell divisions for $N$ generations, leading to $2^N$ daughter cells (Fig. 2). Let $p_m$ be the mutation probability per base per cell division regardless of the type, and assume that the mutations occur independently across sites. Every daughter cell can inherit a subset of substitutions from its parent cell, can mutate a subset of substitutions back to their ancestral states and can generate new substitutions through mutations. The resulting phylogeny is represented as an inhomogeneous Markov process on a binary tree of height $N$. The root represents the mother cell, the leaves represent the $L$ length DNA segments in the $2^N$ daughter cells, and a cell is to its two daughter cells with edges. Let $S_1^1 \ldots S_{2^N}^N$ denote the random number of substitutions in the child segments. The internal nodes in the tree represent the various ancestral forms sampled along the way.

Under these modeling assumptions, the following results ensue. Specific mathematical details are given in **Appendix** section **2.1**. The average substitutions per child for each level $n$ is obtained as $\langle s^{(n)} \rangle = 3p_m L \left[1 + \sum_{a=2}^{n} \prod_{b=a}^{n} \left(\tilde{p}_{inh}^{(b)} - 3p_m\right)\right]$, which is absolutely convergent in $n$. Here $\tilde{p}_{inh}^{(n)} \to .75(1-p_m)$ as $n \to \infty$. For any level $n$ of the tree, the probability distribution over substitutions, $p(S_i^{(n)})$, is identical for all children $i = 1 \ldots 2^n$. Post $N$ generations, when a child is input to the $FST(\delta)$ classifier, it operates with a type I error rate of

$$q_{asexual}^{(N)}(\delta) = p(S_1^{(N)} > \delta L) \tag{1}$$

When processing all children, this error rate leads to $2^N \cdot q_{asexual}^{(N)}(\delta)$ number of type I errors, on average. When the initial population consists of $B_0$ cells *identical* in the the $L$ length DNA region, each undergoing independent exponential growth spurts for $N$ generations, a forest of $B_0$ such binary trees operate, leading to an average number of type I errors of:

$$B_0 \cdot 2^N \cdot q_{asexual}^{(N)}(\delta) \tag{2}$$

Generalization to base-type and position specific mutation rates can be made ( **Appendix** section **2.1.1**) but the qualitative results remain the same: beyond amplification cycles, false classification errors grow with the initial population size.

**Calculations for the Pseudomonas dilution experiment** A single cell dividing once every $\sim 30$ minutes, when grown for $\sim 12$ *hours*, would pass $\sim 24$ generations. For a sequenced 16S hypervariable segment (e.g., V12) length of $250bp$, mutation rate of $\sim 10^{-8}$ per base per cell division, analytic calculations using our cell division model above leads to an average $7.5 \times 10^{-6}$ substitutions in the DNA segment per child ( **Appendix** eqn. (11)). For a 99% sequence similarity threshold, probability of generating an OTU-type child genotype (i.e., deviates by $> \delta L$ substitutions), computed numerically, was: $7.49 \times 10^{-6}$, leading to a total of $\sim 0.0077$ misclassified children. For a 97% sequence similarity threshold, these numbers are essentially equal to zero (less than machine precision).

#### 1.2 Type I error rate of an FST classifier for sequencing molecules post- amplification and sequencing

**Amplification** Amplification via polymerase chain reaction deviates from exponential cell growth in at least one significant fashion. Here, the template DNA strands from the previous generations coexist with newly created strands in every generation, and hence in the final amplification product. So molecules and even their individual strands can vary substantially in the number of extension/mutation cycles they were a product of (**Fig. 3**). This induces varying substitution probabilities across molecules. Results from the previous section are straightforwardly utilized to derive the following results.

Consider a single DNA molecule of length $L$, amplified for $N$ cycles. As before, we restrict to substitution errors alone. We assume mutations are independent across sites, denoting by $p_m$ the per nucleotide mutation probability per base per generation regardless of the type. Each newly generated strand derives its own set of substitutions by inheriting a subset from its template strand and/or newly generating them de novo.

First, assume that the PCR reaction happens with 100% efficiency leading to perfect doubling reactions every cycle. The number of substitutions expected in a template strand that is a result of $n$ extension cycles is the same as that of the previous asexual reproduction case. For all strands that were a product of $n$ extension cycles, the probability distribution over substitutions $\alpha(S_i^{(n)})$ is identical. The type I error rate when classifying a template strand that is a product of $n$ extension cycle is:

$$q_\alpha^{(n)}(\delta) = \alpha(S_1^{(n)} > \delta L)$$

When the FST classifier is input all the strands from the amplification product, the average number of type I errors resulting is $2\sum_{n=1}^{N}\binom{N}{n}q_\alpha^{(n)}$. Here, we have exploited the Moore & Moranas's result [1] that the number of template strands that are a product of $n$ extension cycles is $2\binom{N}{n}$. When there are $B_0$ initial molecules, each undergoing independent perfect doubling reactions for $N$ generations, we obtain:

$$B_0 \cdot 2\sum_{n=1}^{N}\binom{N}{n}q_\alpha^{(n)} \tag{3}$$

type I errors on average. The average number of substitutions observed in the amplification product at generation $m$ is: $\sum_{n=1}^{m}\frac{\binom{m}{n}}{2^n}\langle s^{(n)}\rangle$, with $\langle s^{(n)}\rangle$ defined in the previous section.

When the amplification is imperfect, with imperfection quantified by $\lambda$ ($\lambda = 1$ implying 100% efficiency), the average number of type I errors is given, upto first order, as:

$$B_0 \cdot 2\sum_{n=1}^{N}\lambda^n\binom{N}{n}q_\alpha^{(n)} \tag{4}$$

where we have used Pritchard et al.,'s result that the average number of strands which underwent $n$ extension cycles under imperfect amplification is $2\binom{N}{n}\lambda^n$ [2].

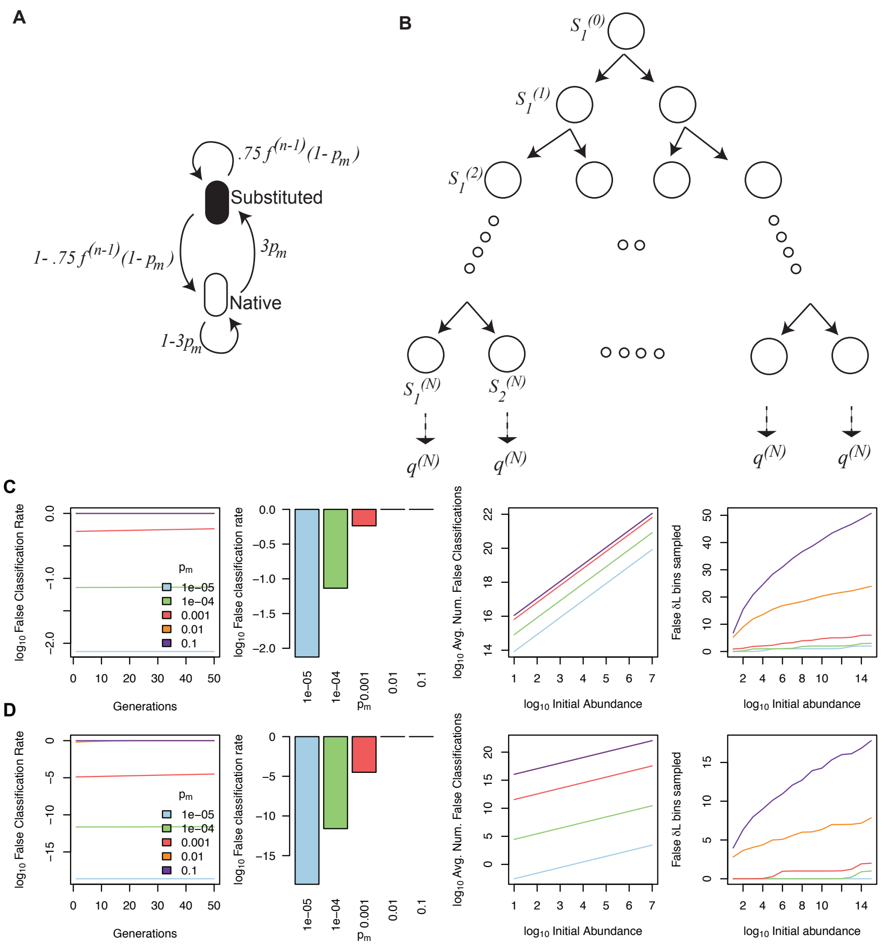


**Figure 2: Cellular reproduction $\rightarrow$ clustering.** Collect the same $L$ length DNA segment from all daughter cells post cell growth for a fixed number of replications $N$, and input to the binary FST classifier as queries, along with the corresponding ancestral mother segment as the reference. We seek the type I error rate of falsely classifying a daughter cell's segment as non-source i.e, not of the mother type. Such non-source type segments go on to generate false OTU varieties. (A) Transition probabilities among substituted and native states of a nucleotide during growth cycle $n$, when the per nucleotide mutation rate is $p_m$. $f^{(n)} \rightarrow 1$ as $n \rightarrow \infty$. (B) Binary tree representing perfect doubling of a cell's DNA segment for $N$ cycles. $S_i^{(n)}$ denotes the realized number of substitutions at daughter cell $i$ in level $n$ of the tree. Each leaf-node on the tree represents one of the final $2^N$ daughter cells' DNA segments resulting from cell growth. $q(N)$ is the probability that a daughter cell (generated naturally) is an OTU-type. $S_i^{(n)}$ evolves according to the process outlined. (C) The false classification rates of an $FST(\delta = .01)$ classifier for various per base per generation mutation rates across generations (left) and at 50 generations (second from left). Average number of artifactual OTU-type daughters (i.e., those that deviate greater than 1% from the mother sequence) due to substitution errors grow with the number of mother cells ("initial abundance") (third from left). Bin sequences that are $k \times (\delta L)$ distance away for $k = 1 \ldots 1/(\delta L)$. Bins contribute to new false microbial discoveries through new sequence clusters. Each bin can form multiple clusters as sequences that are $k \times \delta \times L$ distances apart from the source sequence, can themselves be $\delta L$ distance apart. An increase in number of such bins correspond to an increased number of false microbial discoveries. We plot the expected number of such bins sampled from 99% and 97% similarity threshold based clustering of reads for various per base per generation mutation rates (fourth from left). (D) Same as (C) but for 97% sequence similarity threshold.

**Sequencing** The position-specific quality scores output by the sequencing machine for each sequencing read offers a standard strategy to model sequencing induced substitution probabilities directly. Briefly, quality scores can be transformed to error probabilities conditional on the observed nucleotide, and further integrated to obtain net position-wise substitution probabilities. An inhomogeneous Poisson process parameterized by these probabilities is used to model the total number of substitutions occurring in a sequencing read. The type I error rate is obtained as the probability that the machine induced substitutions exceed the FST threshold as $q_\psi(\delta)$.

$$\psi(S=s) = \int \left[ \sum_{x \in \binom{L}{s}} \prod_{k:x(k)=1} p(k) \prod_{k:x(k)=0} (1-p(k)) \right] dH$$
$$q_\psi(\delta) := \psi(s > \delta L) \tag{5}$$

where $H$ is the distribution function of substitution probability profiles underling sequencing read generation, as obtained from the quality score profiles. The inner sum runs over all possible $L$ length $s$ substituted sequences, $k$ runs over their substituted positions, and $p(k)$ represents the probability of substitution at the site obtained with a suitable transformation from a sequencing machine's quality score profiles.

The expected number of type I errors, when sequencing at a depth of $\tau$ is then:

$$\tau \cdot q_\psi(\delta) \tag{6}$$

**Chaining amplification and sequencing** The aforementioned more exact model for sequencing induced substitutions is cumbersome to model a chain of amplification and sequencing effects. We make a Poisson approximation to the sequencing error model, which models a constant substitution probability across all sites by the estimated average. We found the approximation to have a low error rate ( $\approx 10^{-5}$, based on Serfling [3]) both because of the overall weak per base substitution probabilities by design and because of quality score aware trimming of reads performed by analysts.The latter stabilizes error variance across positions. These simplifications allow a tractable computation of $\psi(s|r)$ (**Appendix** section 2.6), the probability that the sequencing machine outputs an $s$ substituted sequence when input an already $r$ substituted variant from amplification. For the moment, assume amplification efficiencies are comparable across all input sequences and also that there are no further technical biases entering the reaction. Under these conditions, the substitution probabilities of the amplification-sequencing chain are obtained per input molecule as:

$$p(S=s) = \sum_{r=0}^{L} \alpha(r)\psi(s|r)$$
$$q_{\alpha\to\psi}(\delta) := p(S > \delta L). \tag{7}$$

When sequenced at a depth $\tau$, this leads to an expected $\tau \cdot q_{\alpha\to\psi}(\delta)$ number of false classifications.

**Generalizing to a community of input sequences** So far we have restricted ourselves to a contrived system, in which the input to our amplification-sequencing pipeline consisted solely of a single source sequence at some initial abundance. We slightly generalize and consider a community of true input source sequences $i = 1 \dots p$, each with an (apparent) input relative frequency $\eta_i$. It is very reasonable to assume that sequencing machine generates reads in amounts proportional to their input concentrations (see [4, 5]). The relative abundances of source sequences enter the analysis through the following model: $Y_i|\tau \sim Multinomial(\tau, \eta_i)$, where $\tau$ is the sequencing depth and $\eta_i$ is the (apparent) input relative abundance (ref. eqn 2, [5]). Then the net error rate in eqn. (7) can be viewed as a result of integration of input sequence-wise conditionals.

$$q_{\alpha\to\psi}(\delta) = \sum_i \eta_i \cdot q_{\alpha\to\psi}(\delta) := \sum_i q^{(i)}_{\alpha\to\psi}(\delta) \tag{8}$$

Thus, the expected type I errors resulting from each source sequence are predicted as a line along the expected recovered abundance of a source sequence $i$ :

$$q^{(i)}_{\alpha\to\psi}(\delta)\cdot\tau = q_{\alpha\to\psi}(\delta)\cdot\eta_i\cdot\tau = q_{\alpha\to\psi}(\delta)\cdot EY_i \tag{9}$$

Based on eqn. 1 in Kumar et al., 2018 [5], it is straightforward to see varied amplification efficiences across input sequences and the presence of further technical biases alter these equations but the overall qualitative result remains the same: (i) Beyond sample depth, the *apparent* input frequencies (i.e., input frequencies altered by technical biases) is a key determinant of false discoveries. (ii) The false classifications grow as a function of recovered abundance, which upto technical biases, is multiplicative in input abundance and sample depth.

The type I errors made above go on to generate new taxa discoveries algorithmically based on their sequence configurations and abundances. Tracking these microbial discoveries as they continue to aggregate algorithmically from a single source sequence, let alone a mixture of sequences, is difficult. We leave this as an open question, settle for obtaining a rough idea on the scale of such discoveries in **Appendix section 2.2** and proceed to experimentally characterize them in the single colony experiment described in the main text.

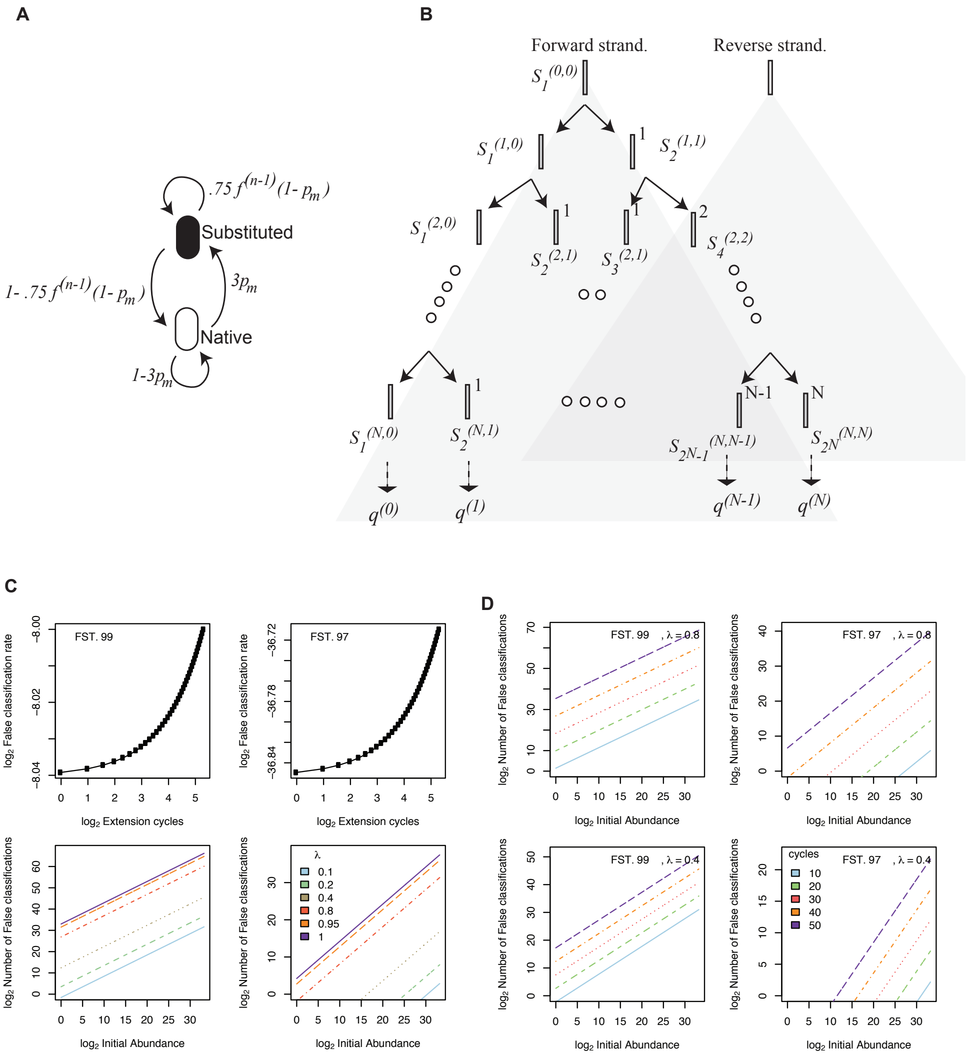


**Figure 3: Amplification $\to$ clustering** (A) Transition probabilities among substituted and native states of a nucleotide during amplification cycle $n$, when the per nucleotide mutation rate is $p_m$. $f^{(n)} \to 1$ as $n \to \infty$. (B) A binary tree representing perfect doubling of a forward strand for $N$ cycles. $S_i^{(n,m)}$ denotes the number of substitutions of a product strand at level $n$ of the tree, while being a product of $m$ amplification cycles. Each leaf-node on the tree represents one of the final $2^N$ child DNA molecules' forward strand product resulting from amplification. $q(N)$ is the probability that a product strand resulting from $N$ cycles of amplification is an OTU-type. $S_i^{(n,m)}$ evolves according to the process outlined. (C) (top row) For two FST threshold settings (99% left column, 97% right column), calculated false classification rate for various extension cycles. Taq polymerase's per nucleotide mutation rate of $1.2 \times 10^{-4}$ per base per doubling, post 40 PCR cycles. (bottom row) Correspondingly, the average number of artifactual OTU-type strands in the amplification product, generated due to substitution errors, grow with the initial concentration ("input abundance"). (D) Rows correspond to two different amplification efficiencies (80% and 40%). In each case, we plot the total number of false classifications. Columns correspond to two different decision rules for the binary classifier: 99% vs 97% sequence similarity. For $\lambda < 1$, we use a first order approximation to calculate the expected behavior of the classifier.

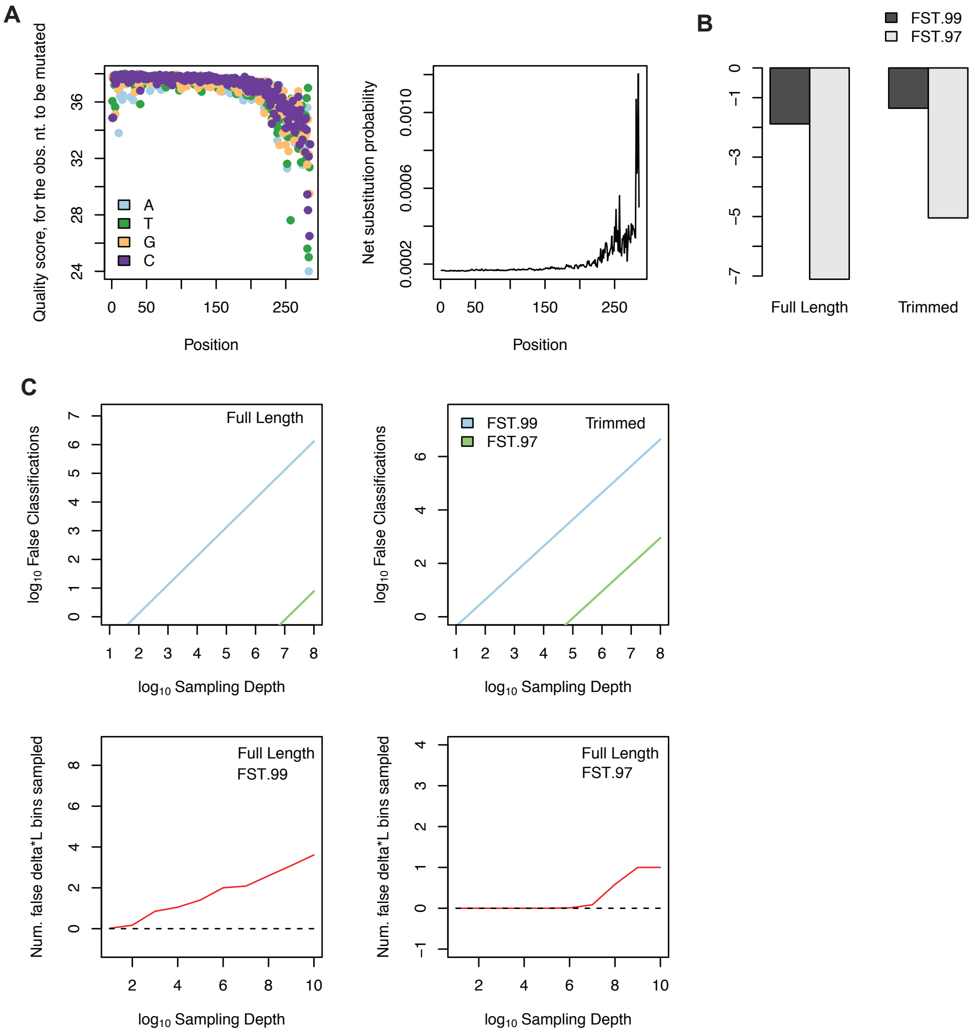


**Figure 4: Sequencing → clustering.** (A) (left) Average quality score profiles derived for the various nucleotide observations and (right) average error probability derived from quality score profiles. (B)Expected false classification rates for 99% and 97% sequence similarity threshold based classifiers, when read trimming is restricted to first 10 bases alone ("full length"), and an additional 85 bases from the end ("trimmed"). The false classification rate is lower for the heavier trim setting. This is because the distance threshold of $\delta L$ is more relaxed (higher). (C) (top row) the expected number of false classifications are visualized as a function of sampled abundance for full length and trimmed settings, at 99% and 97% sequence similarity thresholds. (C) (bottom row) The expected number of false $\delta L$ bins sampled when clustering with 99% and 97% sequence similarity, as a function of sampling depth, for the full length setting.

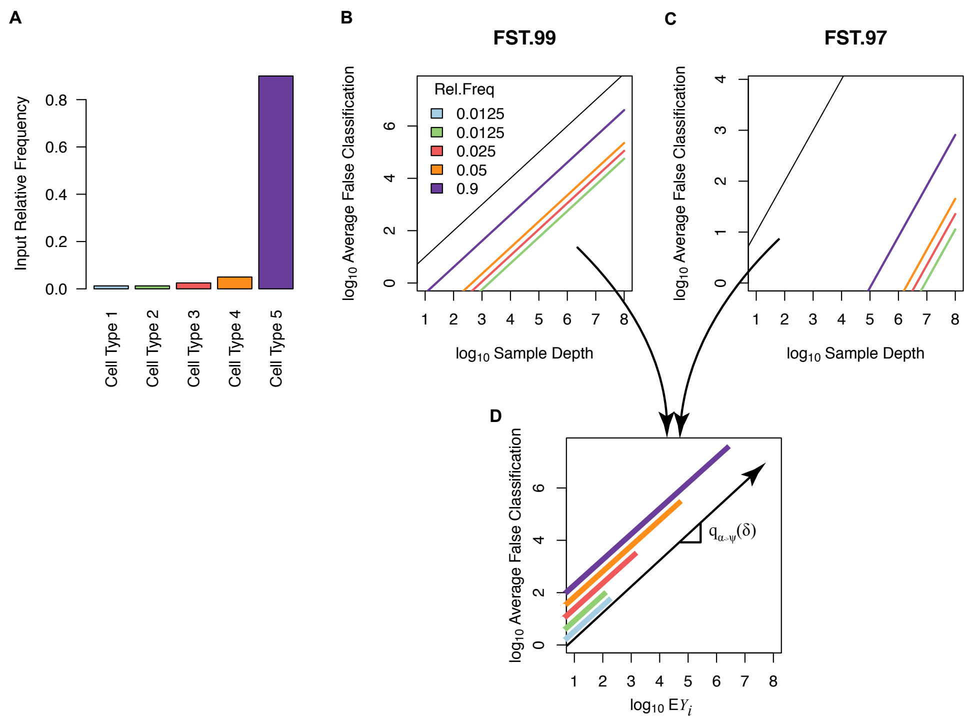


**Figure 5: Amplification→ Sequencing → clustering chain.** Given each source sequence and the set of sequencing reads emitted from a chain of amplification and sequencing processes, the FST classifier decides if the read is source or not based on a fixed sequence similarity threshold. (A)For a community of species with varied input relative frequencies, expected number of false sequencing read classifications are presented as a function of sampling depth for (B) 99% and (C) 97% sequence similarity thresholds respectively. (D) As indicated by the supplementary eqn. (9), the expected number of false classifications for the entire community are captured by two parameters: the expected output abundance of each true source type ($EY_i, i = 1 \ldots 5$ for the five species indicated), and a slope parameter that is the false classification rate, which subsumes the various biochemical and bioinformatics parameters of an amplification-sequencing-clustering chain. The colored bars represent the region along the black line sampled by the colored true input sequences. Amplification and sequencing parameter settings same as in supplementary Figs. 3 and 4.

#### 2 Appendix

##### 2.1 Type I error rate of FST classifiers when genotyping natural variation

**Model** Consider a branching process where a single cell divides asexually for $N$ generations, leading to $2^N$ descendants (**Fig.2B**). Each of the $2^n$ nodes at any level $n$ of the binary tree represents a distinct DNA molecule generated by replicating the ancestral molecule length $L$ for $n$ generations. In addition to assuming perfect doubling, we assume the following: First, we assume mutations occur independently along the entire length of the DNA sequence. Second, we let $p_m$ be the mutation probability for every nucleotide, regardless of its type $\{A,T,G,C\}$. Finally, we shall assume all locations in the sequence suffer the same mutation rate.

At any level $n$, the nodes are indexed left to right on the tree with a subscript $i \in 1,\dots 2^n$ and a superscript $(n)$. Let the random variable $S_i^{(n)} \in \{0,1,\dots L\}$ specify the total number of substitutions generated at node $i$ in level $n$ of the tree. Let $z_i^{(n)}$ indicate the parent node ( in level $n-1$ ) of child node $i$ at level $n$ in the tree. Conditioned on the binary tree, the parent-child relationships $z_i^{(n)}$ is deterministic. For simplicity in notation below, we will be implicit about conditioning on the binary tree (and therefore, $z_\cdot^{(\cdot)}$), $p_m$, $L$ and the source DNA sequence whose amplification is considered.

Consider the process:

$$\begin{aligned} S_1^{(0)} &= 0, \text{ number of substitutions at root node is 0} \\ S_i^{(n)}|X_{i1}^{(n)},X_{i2}^{(n)} &= X_{i1}^{(n)}+X_{i2}^{(n)} \;\forall\; i=1\dots 2^n \text{ nodes, } n=1\dots N \text{ levels} \\ X_{i1}^{(n)}|(S_{z_i^{(n)}}^{(n-1)}=s) &\sim Binomial(s,\tilde{p}_{inh}^{(n)}), \\ X_{i2}^{(n)}|(S_{z_i^{(n)}}^{(n-1)}=s) &\sim Binomial(L-s,3p_m) \end{aligned} \tag{10}$$

In words, the substitution number at the root node, because it corresponds to that of the source sequence, is set to 0. For every non-root node at level $n$, its substitution number $S_i^{(n)}$ is realized by conditioning on the substitution number of its parent node $S_{z_i^{(n)}}^{(n-1)}$ as follows. Given the assumption that the mutations happen independently, the two Binomial random variates $X_{i1}^{(n)}$ and $X_{i2}^{(n)}$ respectively capture the inherited substitutions at a per nucleotide rate of $\tilde{p}_{inh}^{(n)}$ and newly injected substitutions (from non-substituted nucleotides in parent) at a per nucleotide rate of $3p_m$ (**Fig.1A**, and **Appendix 2.3**). The probability of inheritance of a substitution is derived as: $\tilde{p}_{inh}^{(n)} = .75\cdot(1-p_m)\cdot\left[1-(1-4p_m)^{n-1}\right] = .75\cdot(1-p_m)\cdot f^{(n-1)}$ where $f^{(n-1)}\to 1$ as $n\to\infty$. For large $n$, the substitution inheritance probability settles at $.75\cdot(1-p_m)$. The probability of inheritance is a function of the generation number $n$ because per site parental marginals for being a substitution varies with the number of generations passed since the ancestral state.

**Expected substitution numbers** It is interesting to ask for the expected number of substitutions accumulated in each of the $2^n$ product DNA strings after amplification cycle $n$. It is equaly interesting to ask the same question when the amplification itself is driven for a large number of cycles.

The following results appear in **Appendix 2.4**.

(1) The expected number of substitutions is found to be identical for all nodes at a given level $n$, and *scales with* $L$ as:

$$\begin{aligned} E[S_i^{(n)}] &= 3p_m\cdot L\cdot\left[1+\sum_{a=2}^{n}\prod_{b=a}^{n}\left(\tilde{p}_{inh}^{(b)}-3p_m\right)\right] \\ &= E[S_j^{(n)}]\;\forall j=1\dots 2^n] \end{aligned} \tag{11}$$

(2) $\lim_{n\to\infty} E[S_i^{(n)}]$ is well defined and is trapped by:

$$\lim_{n\to\infty} E[S_i^{(n)}] \in \left(3p_mL,\ \frac{3p_mL}{1-\left(\tilde{p}_{inh}^{(n)}-3p_m\right)}\right) \tag{12}$$

**Fig. 6** illustrates the nature of these results for two $p_m$ values. Here the exact averages are calculated numerically and are plotted alongside their upper bound. We see that the average subtitution numbers are strongly dependent on $p_m$. It pays a laboratory to invest in high quality DNA polymerase regardless of the number of amplification cycles run. It is also observed that the average substitution numbers per cycle quickly approach their asymptotic averages. The upper boundary is a rather tight approximation for the expectations.

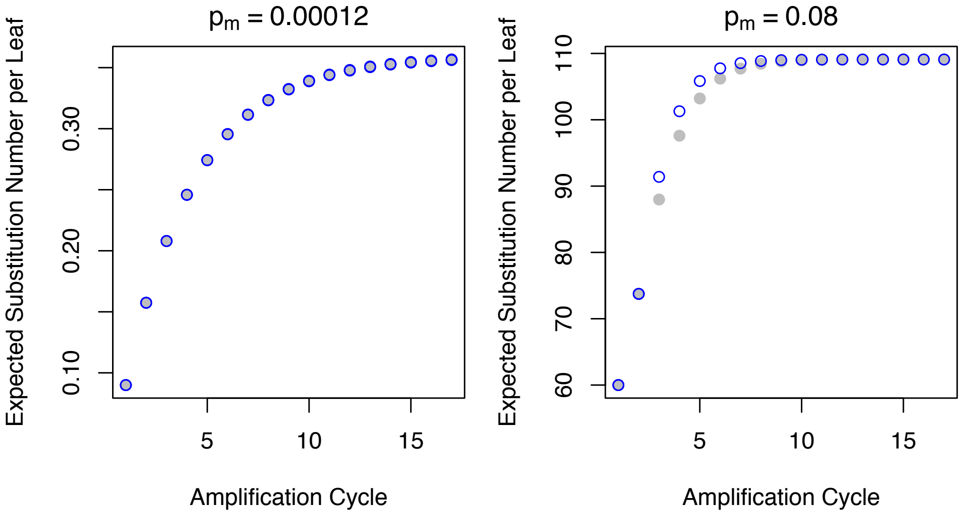


**Figure 6:** Evolution of average substitution numbers is plotted for two per nucleotide mutation rate values ($p_m$). $L = 250$. $p_m = 1.2\times10^{-4}$ is the laboratory measured mutation rate for the *Taq* polymerase. Exact substitution numbers per node (Eqn. (11)) is plotted as filled gray circles as a function of amplification cycle $n$. The respective upper bounds are plotted as open blue circles.

**Probability distribution over substitutions** It is shown that for a given level $n$, the probability distribution over substitutions is *identical* for all nodes at that level (**Appendix 2.5**). This distribution is given as:

$$p(S_i^{(n)} = s_c) = \sum_{s_p=0}^{L} p\left(S_i^{(n)} = s_c | S_{z_i^{(n)}}^{(n-1)} = s_p\right)\cdot p\left(S_{z_i^{(n)}}^{(n-1)} = s_p\right)$$

$$\text{and, } p(S_i^{(n)}) = p(S_j^{(n)}) \text{ for all } j = 1\ldots 2^n, \text{ and for all } n = 1\ldots N \tag{13}$$

The distribution is available numerically given the parameters $p_m$, $L$ and $N$.

**Type I error rate of an FST classifying the children of an ancestral cell** We now answer one of the central questions of this article: Among all the $2^N$ leaf-nodes generated naturally via asexual reproduction, how many are expected to be classifed as non-source type by an FST classifier? It is these classifications that will go on to generate false microbial discoveries as new OTUs. The probability that the $i^{th}$ leaf node is an OTU is the probability that it accumulates a substitution number $> \delta L$. This is given as :

$$q^{(N)}_{asexual} := p(S^{(N)}_i > \delta L) \tag{14}$$

Here the non-dependence of the notation $q^{(N)}_{asexual}$ on $i$ arises because of eqn. (13). Define $T^{(N)}_i$ as the random variable indicating that the leaf node $i$ in level $N$ is a non-source type , and define $T^{(N)}_+ = \sum_{i=1}^{2^N} T^{(N)}_i$. We seek $p(T^{(N)}_+)$.

It is clear $E(T^{(N)}_i = 1) = q^{(N)}_{asexual}$. The expected total number of OTUs is given as:

$$E[T^N_+] = 2^N \cdot q^{(N)}_{asexual} \tag{15}$$

This result is for *per cell* undergoing exponential growth for $N$ generations (i.e., per binary tree with $2^N$ leaf nodes) . If there are $B_0$ identical (identicalness, defined with respect to the $L$ length DNA segment under consideration) copies of the same cell undergoing independent amplifications for the same number of $N$ cycles, there are $B_0$ independent binary trees each generating $2^N q^{(N)}_{asexual}$ OTUs on average, leading to a total of $B_0 \cdot 2^N q^{(N)}_{asexual}$ expected. Thus we conclude that, under the model assumptions, the number of amplification-error induced "artifact" OTUs scale with initial bacterial abundance and amplification cycles. **Fig.1D** shows this behavior for a reasonable choice of parameters.

##### 2.1.1 Generalizing for nucleotide specific mutation rates and position dependencies

The main goal is to relax two of the assumptions made in the simplified model above: a constant mutation rate ($p_m$) per-nucleotide regardless of the nucleotide type, and its location in DNA segment.

The results from the simplified model in the previous subsection is heavily utilized below. As before, the goal is to derive $p(S^{(N)}_i)$ for a given length $L$ sequence.

It is assumed that we know the two dimensional function $p_m(x,y)$, where $x$ indicates the position $x = 1 \dots L$, and $y$ indicates the nucleotide. Fix $N$. For every $x = 1 \dots L$ and for every $y \in \{A,T,G,C\}$, calculate $p(S^{(N)}_i = 1)(x,y)$ from the simplified model in eqn (13) setting $p_m = p_m(x,y)$ and a sequence length of 1 nucleotide. This yields the function $p(S^{(N)}_i)(x,y)$. Consider any two $L$ length sequences $u$ and $v$, and let $v(x)$ retrieve the nucleotide type at position $x$ of sequence $v$. The independence assumption among the sites allows the probability of generating $v$ in a given leaf-node of the $N$ cycle binary tree to be written as:

$$\begin{aligned} p(u \rightarrow v \; in \; a \; given \; leaf-node) &= \prod_{x:\, v(x) \; is \; substituted} p(S^{(N)}_i)(x, v(x)) \prod_{h: v(h) \; is \; not \; substituted} p(S^{(N)}_i)(h, v(h)) \\ \implies \quad p(S^{(N)}_i = s) &= \sum_{i=0}^{\binom{L}{s}} p(u \rightarrow v_i) \end{aligned} \tag{16}$$

where $v_i$ is an $L$ length nucleotide string with $s$ substitutions. This is the *Poisson-Binomial* distribution parameterized by the vector of probabilities $p(S^{(N)}_i)(\cdot, v)$. The OTU generation probability per leaf-node $q^{(N)}_{asexual}$, and the expectation calculations retain the same form as in the simplified model before.

$$\begin{aligned} q^{(N)}_{asexual}(\delta) &:= p(S^N_i > \delta L) \\ E[T^{(N)}_+] &= 2^N \cdot q^{(N)}_{asexual}(\delta) \\ E[\text{OTU type daughters given initial abundance } B_0] &= B_0 \cdot 2^N \cdot q^{(N)}_{asexual}(\delta) \end{aligned} \tag{17}$$

#### 2.2 From type I errors to false microbial discoveries

The type I errors made above go on to generate new taxa discoveries algorithmically based on their sequence configurations and abundances. Tracking these microbial discoveries as they continue to aggregate algorithmically from a single source sequence, let alone a mixture of sequences, is difficult. Consider the following example. There are $\binom{L}{x} \cdot 3^x$ possible nucleotide sequences that are $x$ substitutions away from a given $L$ length source sequence. However, these sequences are not all pairwise $x$ substitutions apart, and hence, experimentally sampled subsets can get clustered in OTU clustering programs. We leave this as an open question and settle for obtaining a rough idea on the scale of such discoveries as a function of similarity thresholds $\delta L$.

Consider an isolated system with a single source sequence and its possible noisy realizations. The distance from the source sequence is binned in units of $\delta L$. The $k^{th}$ bin generates substitutions in the interval $((k-1) \times \delta L, k \times \delta L]$ with probability $p_k = F(k \cdot \delta L) - F((k-1) \cdot \delta L)$. Here $F$ is the distribution function underlying substitutions. For $k = 1$, the left boundary is closed at 0. Sequence falling in the $k = 1$ th bin are subsumed as the source OTU. For the remaining $\delta L$ width bins, both variation in the substitution positions and nucleotides in those positions create new taxa discoveries. We count only a subset of OTU possibilities as follows.

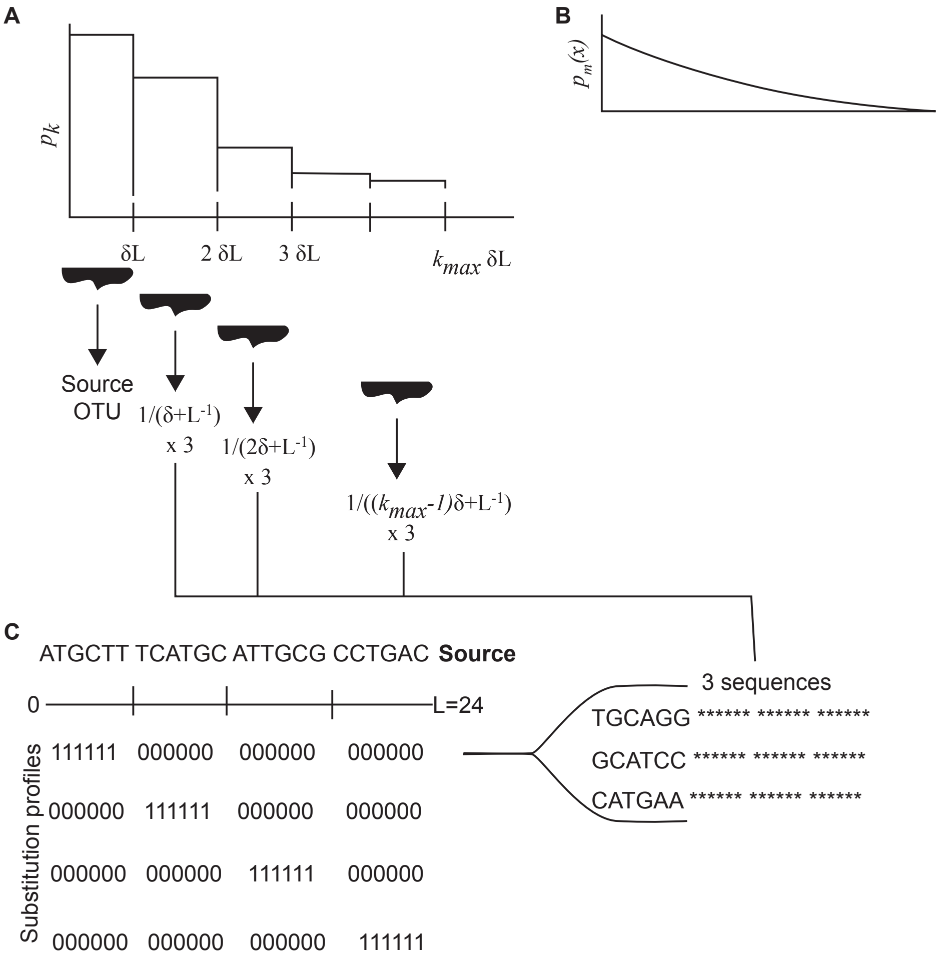


**Figure 7: From type I errors to false microbial discoveries.** The $L$ possible number of substiutions for a length $L$ sequence is binned in $\delta L$ widths. Substituted sequences falling in the first bin is clustered in to the source OTU. The net sampling probabilities for rest of the bins are determined by the cumulative distribution function underlying substitutions, which in turn is determined by the site specific mutation probabilities indicated in (B) as $p_m(x)$, where $x$ is the position. (C) $L$ length substitution profiles with $\delta L$ substitutions in consecutive positions are shown. The total number of such profiles is indicated for each window by the arrows. Each such profile can give rise to three OTU centers $\delta L$ distance away. The total such OTU centers, a subset of all possible OTU types, are obtained by summing along the bins.

First, as also predicted by the simplified models in the previous sections, it is very reasonable to expect the substitution sampling probabilities to fall with the number of subsitutitons, and hence, new OTU centers are likely to be picked from just beyond each bin's left boundary, on average. So consider the total number of substitutions $x \in \{\delta L+1, 2\delta L+1, \ldots, (k_{max}-1)\delta L+1\}$, where $k_{max}$ is an integer chosen such that the probability of generating substitutions beyond $k_{max}\delta L$ is very small. An $L-$ length substitution profile is a bit string that indicates the positions of substitutions with a value 1. For each $x \in \{\delta L+1, 2\delta L+1, \ldots, (k_{max}-1)\delta L+1\}$ , we construct a subset of substitution profiles as follows: divide the source sequence into $\frac{L}{x}$ non-overlapping windows. Correspondingly, generate $L$ length $\frac{L}{x}$ substitution profiles with only the consecutive positions in the $x-$length window holding a value of 1. For any given $x-$ substitution profile, it is straightforward to see that there are 3 nucleotide sequences that are themselves pairwise $x$ substitutions apart. When they occur, they are guaranteed to form distinct OTUs. These are the possible "center" sequences for each $x$. Thus, we have counted about $\frac{L}{x} \cdot 3$ OTUs for $x$ substitutions. As an example, for $x = \delta L + 1$, we have $3 \times \frac{1}{\delta + L^{-1}}$ OTU possibilities.

The total number of OTU possibilities within this restricted set of substitution profiles is then: $\sum_{k=1}^{k_{max}-1} \frac{1}{k\delta + L^{-1}}$. Following Yarza et al., [6] who recommend a $\sim 94.5\%$ sequence similarity clusters for genus reconstructions, we choose $k_{max}(\delta) = .055/\delta$. For $\delta = .01$ (i.e., 99% sequence similarity threshold), $L = 250$, this leads to about 512 artifactual OTUs. For $\delta = .03$ (i.e., 97% sequence similarity threshold), this leads to 90 OTUs. These numbers are one-half to one-third of the total number of taxa found in our experimental study in the next section.

We must note the caveats. We have only considered substitution errors. We have ignored sampling characteristics (not all the aforementioned OTUs would occur in an experiment), ignored further OTU possibilities from substitution profiles with nonconsecutive substitutions within a window, ignored further nucleotide variations within the $k^{th}$ $\delta L$ window that can deviate $\delta L$ away from the anchor strings chosen at $(k-1)\delta L + 1$, and ignored other ensuing complications when there are multiple true input source sequences to consider. Where this analysis can perhaps be reasonably justified is with a single source sequence and when sites have a non-uniform mutation rate. Sort the positions in the decreasing order left to right, such that the probability of substituting is highest on the left end and weakest on the right. On average then, sites accrue mutations left to right. When the probability of non-consecutive substitutions in this sorted space is sufficiently low, ignoring those less likely configurations has a lower impact on these calculations.

#### 2.3 Derivation for substitution ($3p_m$) and substitution inheritance probabilities ($p_{inh}^{(n)}$)

Because there are 4 bases (A, T, G & C), a switch to a new nucleotide (i.e., a substitution) can occur with probability $3p_m$, and a nucleotide remains faithful to itself with probability $1 - 3p_m$. Hence, $X_{i2}^{(n)}$ is determined by a per nucleotide probability of $3p_m$. This result also enforces a natural constraint on $p_m$: $0 \leq 3_pm \leq 1 \implies p_m \in \left[0, \frac{1}{3}\right]$. In this work, we consider the interval $p_m \in \left(0, \frac{1}{3}\right)$ interesting.

On the other hand, the inheritance rate of a substitution as a substitution in the child node is determined both by the nucleotide specification of the substitution in the parent node and the mutation rates determining the inheritance of the same substituted nucleotide / a possible mutation of it to a different nucleotide state that still renders the location in a substituted state in the child node. This is formalized in the derivation below.

For concreteness, consider a single sequence of parent-child relationships represented in the binary tree. We can obtain one such sequence by fixing $i = 1$ in $S_i^{(0)} \to S_i^{(1)} \to S_i^{(2)} ... \to S_i^{(N)}$. So set $i = 1$ until otherwise noted. Let $o_i^{(n,l)}$ indicate the realized nucleotide alphabet at position $l$ in node $i$ on level $n$. Let $\xi_i^{(n,l)}$ indicate if the $l^{th}$ position in child node $i$ at level $n$ is a substitution. Consider the special case that the true nucleotide at the $l^{th}$ position in the original DNA sequence is $o_i^{(0,l)} = A$ (**Fig. 8**). For notation simplicity, we do not explicitly indicate conditioning for $o_i^{(0,l)} = A$ below. The question seeks:

$$\begin{aligned} p(\xi_i^{(n,l)} = 1 | \xi_i^{(n-1,l)} = 1, o_i^{(0,l)} = A) &= \sum_{b \in T,G,C} p(\xi_i^{(n,l)} = 1 | \xi_{z_i^{(n)}}^{(n-1,l)} = 1, o_{z_i^{(n)}}^{(n-1,l)} = b) p(o_{z_i^{(n)}}^{(n-1,l)} = b) \\ &= \sum_{b \in T,G,C} p(\xi_i^{(n,l)} = 1 | \xi_{z_i^{(n)}}^{(n-1,l)} = 1, o_{z_i^{(n)}}^{(n-1,l)} = b) p(o_{z_i^{(n)}}^{(n-1,l)} = b) \end{aligned} \tag{18}$$

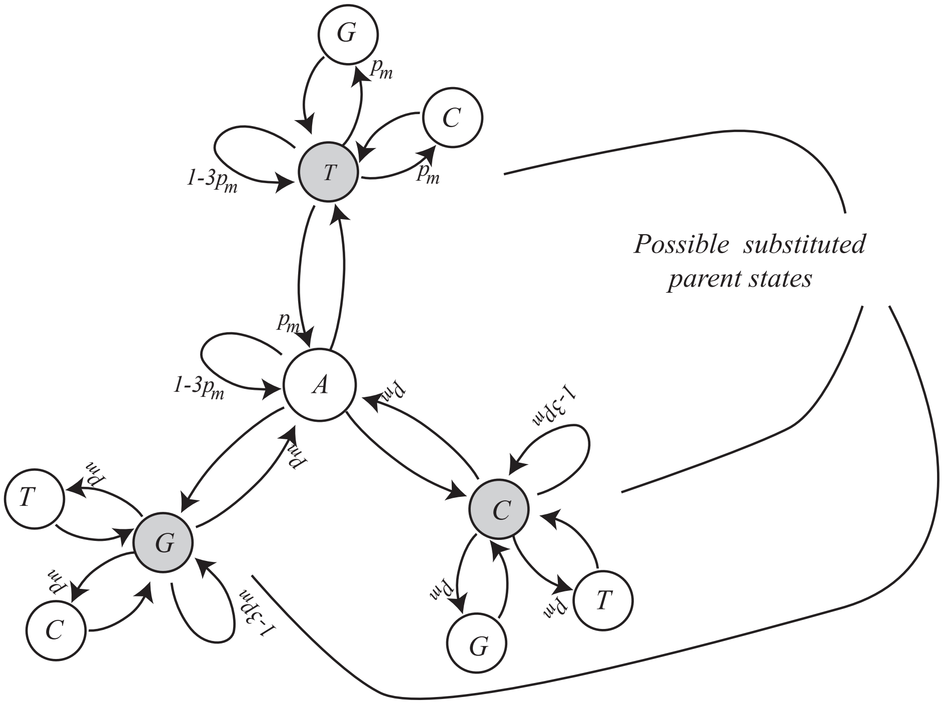


**Figure 8: Possible per-nucleotide substituted states for a parent-node when the true nucleotide state is an** $A$. The transitions among nucleotides that retain a substituted state are shown and their rates identified. Transitions to the true nucleotide state and their probabilities are also described.

Expand each conditional in eqn. 18 as:

$$\begin{aligned} p(\xi_i^{(n,l)} = 1 | \xi_{z_i^{(n)}}^{(n-1,l)} = 1, o_{z_i^{(n)}}^{(n-1,l)} = T) &= p(T \to T) + p(T \to G) + p(T \to C) \\ &= (1-3p_m) + p_m + p_m \\ &= (1-3p_m) + 2p_m \\ &= (1-p_m) \end{aligned} \tag{19}$$

Similarly, obtain:

$$\begin{aligned} p(\xi_i^{(n,l)} = 1 | \xi_{z_i^{(n)}}^{(n-1,l)} = 1, o_{z_i^{(n)}}^{(n-1,l)} = G) = (1-p_m) \\ p(\xi_i^{(n,l)} = 1 | \xi_{z_i^{(n)}}^{(n-1,l)} = 1, o_{z_i^{(n)}}^{(n-1,l)} = C) = (1-p_m) \end{aligned} \tag{20}$$

For the second set of terms in eqn. (18), observe:

$$\begin{aligned} \text{For } b \in \{T,G,C\}, p(o_i^{(n,l)} = b) &= \sum_{b' \in \{A,T,G,C\}} p(o_i^{(n,l)} = b | o_{z_i^{(n)}}^{(n-1,l)} = b') p(o_{z_i^{(n)}}^{(n-1,l)} = b') \\ &= (1-3p_m) p(o_{z_i^{(n)}}^{(n-1,l)} = b) + \sum_{b' \in \{A,T,G,C\} \setminus b} p_m p(o_{z_i^{(n)}}^{(n-1,l)} = b') \\ &= (1-3p_m) p(o_{z_i^{(n)}}^{(n-1,l)} = b) + p_m \sum_{b' \in \{A,T,G,C\} \setminus b} p(o_{z_i^{(n)}}^{(n-1,l)} = b') \\ &= (1-3p_m) p(o_{z_i^{(n)}}^{(n-1,l)} = b) + p_m (1 - p(o_{z_i^{(n)}}^{(n-1,l)} = b)) \\ &= p_m + (1-4p_m) p(o_{z_i^{(n)}}^{(n-1,l)} = b) \end{aligned} \tag{21}$$

Given that $p(o_i^{(0,l)} = A) = 1$ by the particular case we consider here, the following recurrence ensues:

$$
\begin{aligned}
&\text{For } b \in \{T,G,C\}, p(o_i^{(0,l)} = b) = 0 \\
&\qquad p(o_i^{(1,l)} = b) = p_m + (1-4p_m)\cdot 0 = p_m \\
&\qquad p(o_i^{(n,l)} = b) = p_m + \sum_{h=0}^{n-1}(1-4p_m)^h = p_m\left[\frac{1-(1-4p_m)^n}{1-(1-4p_m)}\right] = .25\left[1-(1-4p_m)^n\right]
\end{aligned}
\tag{22}
$$

Applying these results back in eqn. (18), we obtain the inheritance probability:

$$
\begin{aligned}
p(\xi_i^{(n,l)} = 1 | \xi_{z_i^{(n)}}^{(n-1,l)} = 1) &= \sum_{b\in T,G,C} (1-p_m) p(o_{z_i^{(n)}}^{(n-1,l)} = b) \\
&= \sum_{b\in T,G,C} (1-p_m)\cdot .25\left[1-(1-4p_m)^{n-1}\right] \\
&= 3\times(1-p_m)\cdot .25\left[1-(1-4p_m)^{n-1}\right] \\
&= .75\cdot(1-p_m)\cdot \underbrace{\left[1-(1-4p_m)^{n-1}\right]}_{f^{(n-1)}} \\
&:= \tilde{p}_{inh}^{(n)}
\end{aligned}
\tag{23}
$$

where we observe that the inheritance probability is only a function of the level in the tree. The non-dependency of this rate on location, node index $i$ and nucleotide type is made explicit in the notation.

#### 2.4 On expected substitution numbers

Observe:

$$
\begin{aligned}
E[S_i^{(n)} | S_{z_i^{(n)}}^{(n-1)}] &= E[X_{i1}^{(n)} | S_{z_i^{(n)}}^{(n-1)}] + E[X_{i2}^{(n)} | S_{z_i^{(n)}}^{(n-1)}] \\
&= S_{z_i^{(n)}}^{(n-1)} \tilde{p}_{inh}^{(n)} + (L - S_{z_i^{(n)}}^{(n-1)})\cdot 3p_m \\
&= S_{z_i^{(n)}}^{(n-1)}\left[\tilde{p}_{inh}^{(n)} - 3p_m\right] + 3p_m\cdot L \\
\implies E[S_i^{(n)}] &= E[S_{z_i^{(n)}}^{(n-1)}]\cdot\left[\tilde{p}_{inh}^{(n)} - 3p_m\right] + 3p_m\cdot L
\end{aligned}
\tag{24}
$$

As before, we fix $i = 1$, and consider a single sequence of parent-child relationships represented in the binary tree: $S_1^{(0)} \to S_1^{(1)} \to S_1^{(2)} ... \to S_1^{(2^N)}$.

$$\begin{aligned}
E[S_1^{(0)}] &= 0 \\
E[S_1^{(1)}] &= E[E[S_1^{(1)}|S_1^{(n-1)}]] \\
&= E[S_1^{(1)}|S_1^{(n-1)} = 0] \\
&= 3p_m \cdot L \\
E[S_1^{(2)}] &= E[E[S_1^{(2)}|S_1^{(1)}]] \\
&= E[S_1^{(1)}][\tilde{p}_{inh}^{(2)} - 3p_m] + 3p_m \cdot L \\
&= 3p_m \cdot L[\tilde{p}_{inh}^{(2)} - 3p_m] + 3p_m \cdot L \\
E[S_1^{(3)}] &= E[E[S_1^{(3)}|S_1^{(2)}]] \\
&= E[S_1^{(2)}][\tilde{p}_{inh}^{(3)} - 3p_m] + 3p_m \cdot L \\
&= 3p_m \cdot L \cdot \left[1 + \sum_{a=2}^{3} \prod_{b=a}^{3} \left[\tilde{p}_{inh}^{(b)} - 3p_m\right]\right] \\
&\dots \\
E[S_1^{(n)}] &= E[E[S_1^{(n)}|S_1^{(n-2)}]] \\
&= 3p_m \cdot L \cdot \left[1 + \sum_{a=2}^{n} \prod_{b=a}^{n} \left(\tilde{p}_{inh}^{(b)} - 3p_m\right)\right] \\
&\dots
\end{aligned} \tag{25}$$

which is the result indicated in eqn. (11). The result makes it apparent that the right hand side is independent of the child node $i = 1$ for which these calculations were made and is only a function of generation number. Therefore the expected number of substitutions is the same for all $2^n$ nodes at a level $n$, and applies to all sequences of parent-child relationships specified by the binary tree.

To show that $\lim_{n\to\infty} E[S_i^{(n)}]$ exists, first observe that it was derived above that $3p_m$ is the mutation probability for a native nucleotide to switch to a substituted state. So the model defines a natural constraint on $p_m$: $3p_m < 1$ or $p_m < \frac{1}{3}$. Consider the ratio of product terms:

$$\begin{aligned}
\left|\frac{\prod_{b=a}^{n+1}(\tilde{p}_{inh}^{(b)} - 3p_m)}{\prod_{b=a}^{n}(\tilde{p}_{inh}^{(b)} - 3p_m)}\right| &= \left|\tilde{p}_{inh}^{(n+1)} - 3p_m\right| \\
&= \left|.75(1-p_m)(1-(1-4p_m)^{n+1}) - 3p_m\right| \\
&< 1 \text{ for } p_m \in \left(0, \frac{1}{3}\right)
\end{aligned} \tag{26}$$

So the series $\lim_{n\to\infty} \sum_{a=1}^{n} \prod_{b=a}^{n} \left(\tilde{p}_{inh}^{(b)} - 3p_m\right)$ is absolutely convergent. By the comparison lemma for convergent sequences, $\lim_{n\to\infty} \sum_{a=2}^{n} \prod_{b=a}^{n} \left(\tilde{p}_{inh}^{(b)} - 3p_m\right)$ exists as well. So $\lim_{n\to\infty} E[S_i^{(n)}] = \lim_{n\to\infty} 3p_m L \left[1 + \sum_{a=2}^{n} \prod_{b=a}^{n} \left(\tilde{p}_{inh}^{(b)} - 3p_m\right)\right]$ exists.

To derive the bounds in eqn. (12), observe $\tilde{p}_{inh}^{(n)}$ is a monotonically increasing sequence in $n$, bounded above by

$.75(1-p_m)$. So $\left(\tilde{p}_{inh}^{(n)}-3p_m\right) \geq \left(\tilde{p}_{inh}^{(m)}-3p_m\right)$ for all $n > m$. Furthermore, for $p_m \in (0, \frac{1}{3}]$, $|\tilde{p}_{inh}^{(n)}-3p_m| < 1$ for all $n$. So:

$$
\begin{aligned}
3p_m L\left[1+\sum_{a=2}^{n}\prod_{b=a}^{n}\left(\tilde{p}_{inh}^{(b)}-3p_m\right)\right] &\leq 3p_m L\left[1+\sum_{a=2}^{n}\prod_{b=a}^{n}\left(\tilde{p}_{inh}^{(n)}-3p_m\right)\right] \\
&= 3p_m L\left[1+\sum_{a=2}^{n}\left(\tilde{p}_{inh}^{(n)}-3p_m\right)^{n-a+1}\right] \\
&= 3p_m L\left[\sum_{a=0}^{n-1}\left(\tilde{p}_{inh}^{(n)}-3p_m\right)^{a}\right] \\
&= 3p_m L\left(\frac{1-\left(\tilde{p}_{inh}^{(n)}-3p_m\right)^{n}}{1-\left(\tilde{p}_{inh}^{(n)}-3p_m\right)}\right)
\end{aligned}
\tag{27}
$$

#### 2.5 On the probability distribution over substitution numbers

Write the conditional probabilities in eqn. (10) equivalently as:

$$
\begin{aligned}
S_1^{(0)} &= 0, \text{ number of substitutions at root node is 0} \\
p\left(S_i^{(n)} = s_c | S_{z_i^{(n)}}^{(n-1)} = s_p\right) &= \sum_{k=0}^{s_c} Binomial(k|s_p, \tilde{p}_{inh}^{(n)}) \cdot Binomial(s_c - k|L - s_p, 3p_m) := \Gamma(s_c, s_p; n)
\end{aligned}
\tag{28}
$$

With eqn. (28), numerically compute an $(L+1)\times(L+1)$ conditional probability matrix $\Gamma$ for every amplification cycle $n$. Similarly, conditioning on the substitutions of the parent node, the child node's substitutions are conditionally independent of the rest of the nodes on the tree. So for any non-root node $i$ at any level $n$, write:

$$
\begin{aligned}
p\left(S_i^{(n)} = s_c\right) &= \sum_{s_p=0}^{L} p\left(S_i^{(n)} = s_c | S_{z_i^{(n)}}^{(n-1)} = s_p\right) \cdot p\left(S_{z_i^{(n)}}^{(n-1)} = s_p\right) \\
&= \sum_{s_p=0}^{L} \Gamma(s_c, s_p; n) \cdot p\left(S_{z_i^{(n)}}^{(n-1)} = s_p\right)
\end{aligned}
\tag{29}
$$

With eqn. (29), numerically compute $p(S_i^{(n)})$ for any level $n$.

To show that $p(S_i^{(n)}) = p(S_j^{(n)})$ for all $j = 1\ldots 2^n$, and for all $n$, a proof by induction follows. The statement holds for $n = 1$.

$$
\begin{aligned}
p(S_1^{(0)}) &= 0 \text{ by construction.} \\
p(S_1^{(1)} = s_c) &= \sum_{s_p=0}^{L} p(S_1^{(1)} = s_c | S_1^{(0)} = s_p) p(S_1^{(0)} = s_p) \\
&= p(S_1^{(1)} = s_c | S_1^{(0)} = 0) \\
&= \Gamma(0, s_p; 1) \\
&= p(S_2^{(1)} = s_c | S_1^{(0)} = 0) \\
&= \sum_{s_p=0}^{L} p(S_2^{(1)} = s_c | S_1^{(0)} = s_p) p(S_1^{(0)} = s_p) \\
&= p(S_2^{(1)} = s_c)
\end{aligned}
\tag{30}
$$

Assuming true for level $n$, it is shown that the statement holds for level $n+1$.

$$\begin{aligned}
p(S_1^{(n+1)} = s_c) &= \sum_{s_p=0}^{L} p(S_1^{(n+1)} = s_c | S^{(n)}_{z_1^{(n+1)}} = s_p) p(S^{(n)}_{z_1^{(n+1)}} = s_p) \\
&= \sum_{s_p=0}^{L} \Gamma(s_c, s_p; n+1) p(S^{(n)}_{z_1^{(n+1)}} = s_p) \\
&= \sum_{s_p=0}^{L} \Gamma(s_c, s_p; n+1) p(S^{(n)}_{z_j^{(n+1)}} = s_p) \ \ \forall\, j = 2 \ldots 2^{n+1} \\
&= \sum_{s_p=0}^{L} p(S_j^{(n+1)} = s_c | S^{(n)}_{z_j^{(n+1)}} = s_p) p(S^{(n)}_{z_j^{(n+1)}} = s_p) \\
&= p(S_j^{(n+1)} = s_c)
\end{aligned} \tag{31}$$

#### 2.6 $\psi(s|r)$ computation

For each sequencing read $i$, its position-specific Phred scores $q_i(k)$, over positions $k = 1 \ldots L$ specify the conditional probability that the observed nucleotide at the position $k$ is an error through the transformation $10^{-q_i(k)/10}$. Following the approximation noted in the subsection "Chaining amplification and sequencing", we calculate the average error rate for each read, $\bar{\lambda}_i = L \cdot \frac{1}{L} \sum_{k=1}^{L} 10^{-q_i(k)/10}$ and denote by $\hat{f}_\lambda$, their empirical density across reads. We can then numerically compute:

$$\psi(S = s | r = 0) := \int_t p(s|t) \hat{f}_\lambda(t) dt \tag{32}$$

where $p(s|t) = Poisson(t)$. Numerical calculations of other quantities like $q_\psi(\delta)$ follow in a straightforward fashion.

Calculation of $\psi(S|r)$ for all $r > 0$ follow in exactly the same fashion as eqn. (10) where the Binomials are replaced by the corresponding Poissons.

For computational tractability, we do these calculations over a random subset of $10,000$ sequencing reads.

### Supplementary Note 2

#### Supplementary Note 2: The dynamics of false species accumulations in a population survey

Here we study the accumulation dynamics of false species types, generated through misclassification of true species types in a population survey. We incorporate false species types within the sampling theoretic framework of Chao [1] and Harris [2] and study their accumulation behavior analytically and with simulations. An interpretation for the Prokounter's linear predictor proposed in the main manuscript is offered by placing it in the context of observed and asymptotic richness estimates.

Throughout, both the number of true species and the possible false varieties are assumed finite. Thus, we let $i = 1 \ldots I$ denote the true number of species with population relative frequencies $p_i$. Every true species $i$ can be falsely classified as one of $j = 1 \ldots J$ species with probabilities $\eta_i q_{ij}$, where $\eta_i$ is the probability of misclassifying species type $i$ and $q_{ij}$ represents the conditional probability that the misclassification is to a false type $j$. Then the the probability of sampling a true type $i$ is $p_i(1-\eta_i)$ and the probability of sampling a false type $j$ is $\sum_{i=1}^{I} p_i \eta_i q_{ij}$.

**Case 1** We first assume that none of the false species types are identical to any of the true species types. Following Chao [1], we consider the random variable $n_r$, which is the frequency of frequency $r$ in the observed count data i.e., the number of species registering a count of $r$. We decompose $n_r$ as a sum of two terms: $n_r = n_r^{(T)} + n_r^{(F)}$, where $n_r^{(T)}$, corresponds to the number of observed true species with frequency $r$ and $n_r^{(F)}$ corresponds to the observed number of false types with frequency $r$. Let the sampling depth be $N = \sum_r r n_r$. Then, based on eqn. 2 in [1], the average value of $n_r$ specified as a function of sampling depth $N$ can be decomposed as:

$$En_r(N) \approx \underbrace{\sum_{i=1}^{I} \frac{(Np_i(1-\eta_i))^r e^{-Np_i(1-\eta_i)}}{r!}}_{:=\text{expected true species with frequency } r,\ En_r^{(T)}(N)} + \underbrace{\sum_{j=1}^{J} \frac{(N\sum_i p_i\eta_i q_{ij})^r e^{-N\sum_i p_i \eta_i q_{ij}}}{r!}}_{:=\text{expected false types with frequency } r,\ En_r^{(F)}(N)} \tag{1}$$

The total possible false species types $J$ is appropriately recovered at $N = 0$ and $r = 0$ :

$$En_0^{(F)}(N=0) = \sum_{j=1}^{J} e^{-N\sum_i p_i \eta_i q_{ij}}|_{N=0} = J. \tag{2}$$

Let $\tilde{J}(N)$ denote the total number of false species types observed when sampling at a depth of $N$. Thus the expected false species accumulation $E\tilde{J}(N)$ is:

$$\begin{aligned} E\tilde{J}(N) = &\underbrace{En_0^F(0)}_{\text{total false types possible}} - \underbrace{En_0^F(N)}_{\text{average number of as yet undiscovered false types}} \\ &= J - \sum_{j=1}^{J} e^{-N\sum_i p_i \eta_i q_{ij}} \end{aligned} \tag{3}$$

For a given sampling effort $N$, the total apparent richness (I+J) is decomposed into four categories: number of observed true types ($\tilde{I}(N)$), number of observed false types ($\tilde{J}(N)$), number of undetected true types ($\tilde{I}_u(N)$), number of undetected false types ($\tilde{J}_u(N)$). It is clear from eqn. (3), $E\tilde{J}(N) \to J$ as $N \to \infty$ appropriately indicating that, with sufficient sampling, all possible false discoveries are recovered. Similarly, it can be shown $E\tilde{J}_u(N) \to 0$, $E\tilde{I}(N) \to I$, and

$E\tilde{I}_u(N) \to 0$ as $N \to \infty$.Then the average values of observed richness, $S^{obs}$, follow our intuition:

$$ES^{obs}(N) = E\tilde{I}(N) + E\tilde{J}(N) \to I + J, N \to \infty \tag{4}$$

Thus, without appropriately accounting for false discovery generation, any asymptotically consistent total richness estimator, $\hat{S}(N)$, estimates the total apparent richness of the community as $I+J$, overestimating its true richness.

$$E\hat{S}(N) = E\tilde{I}(N) + E\tilde{J}(N) + E\tilde{I}_u(N) + E\tilde{J}_u(N) + B(N) = I + J + B(N) \to I + J, N \to \infty. \tag{5}$$

where $B(N) \to 0, N \to \infty$ is a bias term.

The Chao1 estimator uses the observed richness $S^{obs}(N)$ to estimate $E\tilde{I}(N)$ and estimates a lower bound for undetected richness with $\frac{En_1^2}{2En_2}$. The two terms are biased by false discoveries as below:

$$\begin{aligned} E\left(S^{obs}(N) - E\tilde{I}(N)\right) &= E\tilde{J}(N) \\ \frac{En_1^2(N)}{2En_2(N)} &= \frac{\left(En_1^{(T)}(N) + En_1^{(F)}(N)\right)^2}{2\left(En_2^{(T)}(N) + En_2^{(F)}(N)\right)} \end{aligned} \tag{6}$$

Appropriate bias correction factors are therefore needed.

Fig. 1 provides a multinomial simulation illustration of the problem setup for various sampling effort $N$. In the presence of false discoveries, observed and asymptotic richness estimates far exceed true richness and steadily approach the apparent asymptotic richness of the community $I+J$. Furthermore, the sampling depth needed to cover the true community is much lower than what the survey data including false discoveries would indicate.

**Case 2** We now allow for the general possibility that true types can also be misclassified to a subset of the true species types. In this case, we decompose the true species types in to two further sub-types: true types that occur because of correct classification and true types that occur because of misclassification from another true type. If $J^*$ represents the number of false types that are not identical to any of the true types, then the total false classifications that could arise are $J = J^* + I$. Eqn. (1) gets adjusted to the following and similar discussions as in Case 1 follow.

$$En_r(N) \approx \underbrace{\sum_{i=1}^{I} \frac{(Np_i(1-\eta_i))^r e^{-Np_i(1-\eta_i)}}{r!}}_{\text{expected true types with freq } r, En_r^{(T)}(N)} + \underbrace{\underbrace{\sum_{i=1}^{I} \frac{(N\sum_{k:k\neq i} p_k \eta_k q_{ki})^r e^{-N\sum_{k:k\neq i} p_k \eta_k q_{ki}}}{r!}}_{\text{expected number of true types because of misclassification at freq. } r} + \sum_{j=1}^{J^*} \frac{(N\sum_i p_i \eta_i q_{ij})^r e^{-N\sum_i p_i \eta_i q_{ij}}}{r!}}_{\text{expected false types at freq. } r, En_r^{(F)}(N)} \tag{7}$$

**Case 3** We consider a further simplification to case 1 above. Specifically, we shall assume $\eta_i = \eta$, and all $q_{ij} = q$. Eqn. (1) gets adjusted to:

$$En_r(N) \approx \underbrace{\sum_{i=1}^{I} \frac{(Np_i(1-\eta))^r e^{-Np_i(1-\eta)}}{r!}}_{\text{expected true discoveries with freq. } r,\ En_r^{(T)}(N)} + \underbrace{J\frac{(N\eta q)^r e^{-N\eta q}}{r!}}_{\text{expected false types with freq. } r,\ En_r^{(F)}(N)} \tag{8}$$

Correspondingly, the expected number of false discoveries for a prescribed sampling effort $N$, from eqn. 3 gets adjusted to:

$$E\tilde{J}(N) = J - Je^{-N\eta q} = J \cdot (1 - e^{-N\lambda}) \tag{9}$$

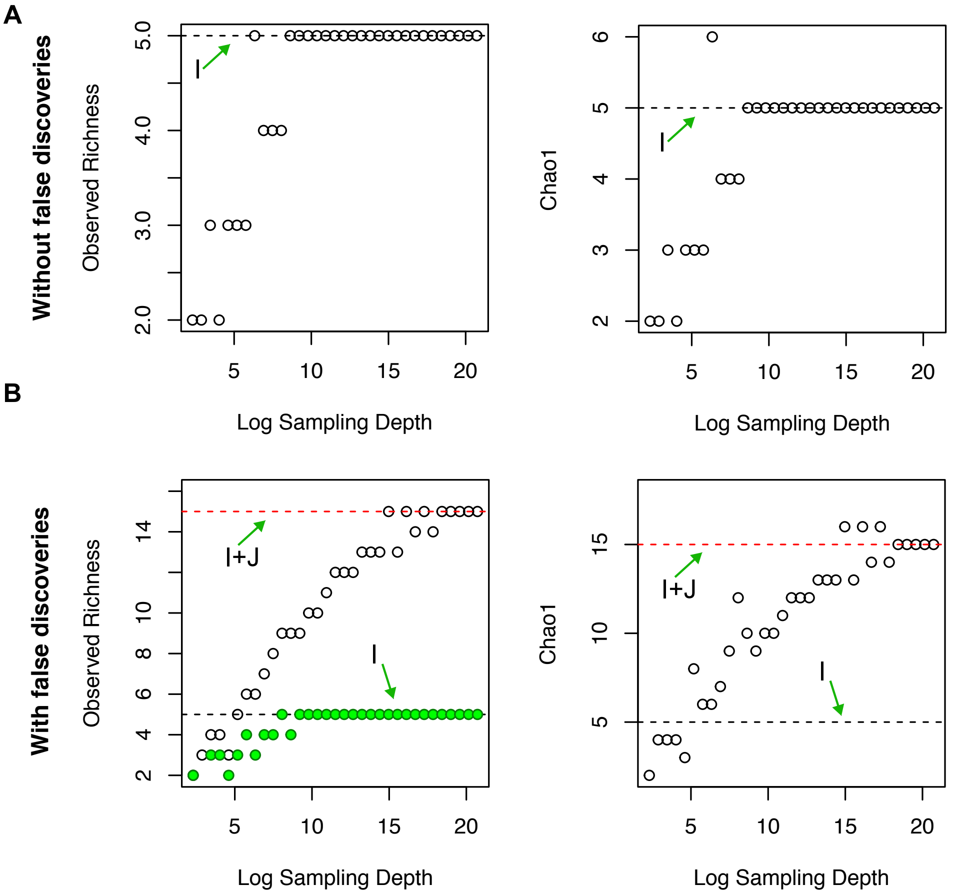


**Figure 1:** (A) We construct an artificial in silico community with $I = 5$ true types with $p_i \propto \frac{1}{.1^i \cdot i}$, for $i = 1 \ldots 5$. In (B) each of the 5 true types can also be misclassified into to two possible false types with conditional probabilities $q_{i\cdot} \propto (1, 1/2)$ for a total of $J = 10$ false types. Thus the apparent asymptotic richness is $I + J = 15$. Other parameters: $\eta_i = \eta = .01$. Points in green indicate the sampled true richness of the community (i.e., only true species) and the black points indicate the observed apparent richness (true + false species) of the community. In the presence of false discoveries, with sufficient sampling, both the observed and traditional asymptotic richness estimator (here Chao1), overestimate true richness. Observed and asymptotic richness estimates correlate well until the total apparent richness is sufficiently sampled.

.

where $\lambda = \eta q$ is an effective parameter.

Fig. 1 motivates the following idea, which is illustrated in Fig. 2. Let $N^*$ denote the sampling effort needed to recover all true species with high probability. Let $N^{**}$ denote the sampling effort needed to recover true *and* all false species types with high probability. With sequencing technologies, it is very likely $N^{**} > N^*$. The sampling effort axis is then split into three windows: $w_1 = [0, N^*]$, $w_2 = (N^*, N^{**})$, $w_3 = [N^{**}, \infty)$.

In the window $w_2$, the gradient of the expected observed number of features is, with high probability:

$$\frac{dES^{obs}(N)}{dN} = \frac{d\left(E\tilde{I}(N) + E\tilde{J}(N)\right)}{dN} \approx \frac{d(I + E\tilde{J}(N)}{dN} = J \cdot \lambda e^{-N\lambda} \tag{10}$$

from eqn. (9). Thus within window $w_2$,

$$\log \frac{dES^{obs}(N)}{dN} \approx \underbrace{(\log J + \log \lambda)}_{\text{intercept}} \underbrace{-\lambda}_{\text{slope}} N \tag{11}$$

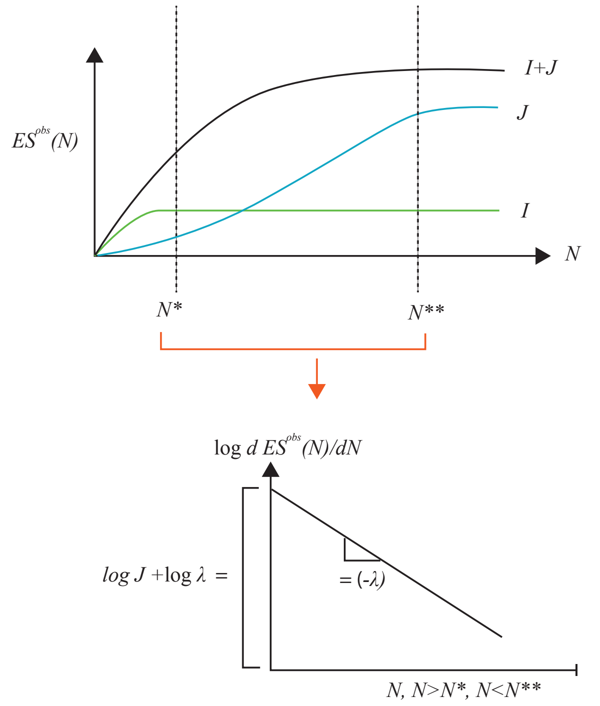


**Figure 2:** In the simplified case 3, regressing logged species accumulation rate against sampling effort in the sampling regime $w_2 = (N^*, N^{**})$, yield estimates of asymptotic false richness generated from a survey and the effective species mislassification rate $\lambda$.

.

with high probablity. Then, under this simplified model, a trend line of the log gradient of the expected observed species richness (a rasonable stimate is the log gradient of species rarefaction curve for various $N \in w_2$) against sampling effort $N$ can be used to calculate both the total number of possible false discoveries $J$, and the net false classification rate $\lambda$. Knowledge of these quantities can allow us to establish appropriate corrections for asymptotic richenss estimates and also for quantities that underly species richness calculations.

The problem reduces to identifying $w_2$, which involves two parameters $N^*$ and $N^{**}$. If one assumes that practical sampling limitations does not allow us to ever reach $N^{**}$, $N^*$ alone needs to be estimated. We leave this as an open question and settle for varying $N^*$ through all values $N$ and getting a sense of the range of $\lambda$ and $J$ values that arise.

##### Prokounter's proposal

In principle, a finite sampling correction for the observed richness and asymptotic richness estimates is achieved with access to an unbiased estimator $\widehat{E\tilde{J}(N)}$ for $E\tilde{J}(N)$as:

$$\begin{aligned} \widehat{S^{obs}(N)} &= S^{obs}(N) - \widehat{E\tilde{J}(N)};\ E\widehat{S^{obs}(N)} \to I, N \to \infty \\ \widehat{\hat{S}(N)} &= \hat{S}(N) - B(N) - J \approx \hat{S}(N) - B(N) - \widehat{E\tilde{J}(N)};\ E\widehat{\hat{S}(N)} \to I, N \to \infty. \end{aligned} \tag{12}$$

However, in both cases 1 and 2 above, $E\tilde{J}(N)$ is defined based on a number of unknown parameters. In a small survey like a supervised local bird walk where an expert accompanies amateur birders, perhaps $I$ and $J$ are small enough that direct estimation of the average rate parameters are feasible. In large scale automated surveys like 16S surveys, estimation is a challenging problem. The methodology in the main manuscript proposes to circumvent this problem specifically in 16S surveys through a function $f_t(\log N)$ estimated from within-genus taxa accumulation data. Specifically, the methodology assumes that most within-genus taxa discoveries across detected genera are false. In terms of notation, the main manuscript has the following correspondence for within-genus richness analysis: $N \equiv y_{gj}$, $S^{obs} \equiv n_{gj}$. The linear predictor in the main text models:

$$\begin{aligned}
\log ES^{obs}(N) &= \log E\left[\tilde{I}(N) + \tilde{J}(N)\right] \\
&= \log E\left[I - \tilde{I}_u(N) + \tilde{J}(N)\right] \\
&= \log\left[I \underbrace{-E\tilde{I}_u(N))}_{\text{sampling effort bias}} + \underbrace{E\tilde{J}(N)}_{\text{false discovery bias}}\right] \\
&\approx \underbrace{X\beta}_{\text{design variables}} + f_t(\log N)
\end{aligned} \tag{13}$$

where adjustments for false discovery accumulation and sampling effort $N$ are folded into $f_t(\cdot)$ and any additional predictors we may wish to incorporate. When asymptotic richness estimates are used as response variables, a similar interpretation follows involving the bias term $B(N)$ in eqn. (12).

### Supplementary Note 3

#### Supplementary Note 3 : Simulations

To illustrate that skewed differential abundance of genera can lead to confounded sample-wide differential richness inference with current estimators, we built the following two-group simulator. The number of noisy taxa generated for a given genus is derived as a function of genus abundance .

The simulation procedure is executed independently for each sample-group, and genus category. We therefore do not explicitly indicate the genus and sample group in the notation below. $j$ indexes observations, $i$ indicates the true types within the genus, $k$ indexes the set of taxa (artifactual + 1 true type ) generated from the true type. The recovered abundance for a taxon is indicated as $y_{jik}$. $+$ in a subscript index indicates summing over that index.

$$\begin{aligned}
& y_{jik}|y_{ji+}, \theta_{i\cdot} \sim Multinomial(y_{ji+}, \theta_{ji}) \\
& \theta_{ji} \sim Dirichlet(1, \frac{1}{2}, \frac{1}{3}, \cdots \frac{1}{\tilde{n}_{ji}}) \\
& \tilde{n}_{ji}|y_{ji+} = \lceil e^{f_t(y_{ji+})} \rceil \\
& y_{ji+}|y_{j++}, \nu_{j\cdot} \sim Multinomial(y_{j++}, \nu_{j\cdot}), \nu_{ji} \propto u_{ji} \\
& u_{ji} \sim Uniform(0,1), i = 1 \ldots n_j^0
\end{aligned} \tag{1}$$

In words, a given genus-wide recovered abundance $y_{j++}$ is distributed among its constituent biological types (species) as $y_{ji+}$. Each species-specific abundance is further distributed among all the taxa (true + artifactual taxa) constituting the species as $y_{jik}$ for $k = 1 \ldots \tilde{n}_i$. The number of taxa generated for a true species type $\tilde{n}_{ji}$ is a function of its recovered abundance $y_{ji+}$ for the species. The Dirichlet parameter is set such that on average, the first category is the dominant, corresponding to the true taxon for the type. The rest are artifactual. For the null case, the true richness $n^0$ are input parameters, are not varied for samples within a group. Genus-wide recovered abundances $y_{j++}$ are sampled from any given real dataset. The median genus-wide recovered abundances were used as a threshold to define "low" and "high" values for inducing differential abundance. Additional Guassian noise is injected in the sampled recovered abundance values. $f_t(\cdot)$ is a fitted loess trend for within-genus taxa accumulation data.

### Supplementary Note 4

#### Supplementary Note 4: SSANOVA model construction

Here, we present more details on the construction of the smoothing spline ANOVA model specified in equation (1) of the main text. We assumed for $z_{gj} = \log n_{gj}$:

$$E[z_{gj}|g, y_{gj}] = \eta(g, y_{gj}) + \varepsilon_{gj} = \kappa + f_R(\log y_{gj}) + f_{GR}(g, \log y_{gj}) + \varepsilon_{gj},\ \varepsilon_{gj} \sim N(0, \sigma^2)$$

Here, $\eta$ is assumed a member of a reproducing kernel Hilbert space (RKHS) of functions $H$ with the orthogonal decomposition $H = H_0 \oplus H_1$. $H_0$ is a finite dimensional space spanned by a set of basis functions $\phi_1(\cdot), \ldots, \phi_m(\cdot)$. $H_1$ is an RKHS with an associated bivariate, symmetric reproducing kernel $R_1(\cdot,\cdot)$. Suppose we wish to model a given set of responses $\tilde{y}_1, \ldots, \tilde{y}_n$, through a smooth function $\eta$ over predictors $x_1, \ldots, x_d$. It can be shown that $\eta(x_i)$ is conveniently decomposed orthogonally as $\eta(\cdot) = \eta_0(\cdot) + \eta_1(\cdot) + \rho(\cdot)$, with $\eta_0(\cdot) \in Span(\{\phi_i, i = 1 \ldots m\})$, $\eta_1(\cdot) \in Span(\{R_1(x_i, \cdot), i = 1 \ldots n\})$ and $\rho(\cdot) \in H_1 \ominus Span(\{R_1(x_i, \cdot), i = 1 \ldots n\})$.

For estimating $\eta$, consider the penalized optimization problem:

$$\hat{\eta} = \arg\min_{h \in H} l(h|\tilde{y}_., x_.) + \lambda J(h), \tag{E1}$$

where $l(\cdot|\tilde{y}_., x_.)$ is the negative log likelihood, $J(\cdot)$ is a roughness penalty, and $\lambda$ a regularization parameter. $J(h)$ is a squared semi-norm in $H_1$ and penalizes overfitting $h$ to data in $H_1$. The celebrated Kimeldorf-Wahba[1] theorem proves that the solution to this penalized optimization problem has a finite dimensional representation:

$$\hat{\eta}(\cdot) = \sum_{i=1}^{m} c_i \phi_i(\cdot) + \sum_{i=1}^{n} r_i R_1(x_i, \cdot) \tag{E2}$$

This allows one to write $J(h) = r_.^T R_1 r_.$ , where the $(i, j)^{th}$ element of $R_1$ is $R_1(x_i, x_j)$. As an example, a cubic smoothing spline on $[0, 1]$ is obtained as solution to the following instance of the penalized optimization problem in eqn. (E1): $\tilde{y}_i = \eta(x_i) + \varepsilon_i, \varepsilon_i \sim N(0, \tau^2), x_i \in [0, 1], J(\eta) = \int_0^1 \left(\frac{d^2\eta}{dx^2}\right)^2 dx$.

Construction of the recovered abundance dependent model in eqn. (1) in the main text proceeds as follows. As per example 2.7 in Gu[2], we specify marginal reproducing kernels for (A) the discrete space of genera $\{1 \ldots K\}$, where $K$ denotes the number of detected genera, and (B) the continuous space of $\log y_{gj}$ rescaled to $[0, 1]$ via the transformation $t_{gj} = \frac{y_{gj} - max(y_{..})}{max(\log y_{..}) min(\log y_{..})}$. Reproducing kernels for the product space $\{1 \ldots K\} \otimes [0, 1]$ - decomposed in to orthogonal subspaces corresponding to the intercept, genus-specific main effects term, (rescaled) recovered abundance dependent main effects term, and the interaction terms - are constructed as tensor products of marginal reproducing kernels. These are derived in example 2.7 and presented in table 2.5, Gu[2]. The intercept and any linear dependence on the $t_{gj}$ are members of $H_0$, spanned by the basis $\{\phi_1(x) = 1, \phi_2(x) = x - 0.5\}$, $x \in [0, 1]$. The higher order terms are penalized for roughness and are elements of $H_1$, the reproducing kernel $R_1(\cdot,\cdot)$ of which is constructed using table 2.5, Gu[2]. Specifications of $\phi_.(\cdot)$ and $R_1(\cdot,\cdot)$ then enter the optimization problem in eqn. (E1) where $h$ has the form specified in eqn. (E2). To estimate $\kappa$, $f_R(\cdot)$ and $f_{GR}(g, \log y_{gj})$ using the `gss` package, we pass the R model formula $z_{gj} \sim\ g * log(y_{gj})$ to the `gss::ssanova` function.

[2] Chong Gu. *Smoothing spline ANOVA models*, volume 297. Springer Science & Business Media, 2013.

### Supplementary Figures

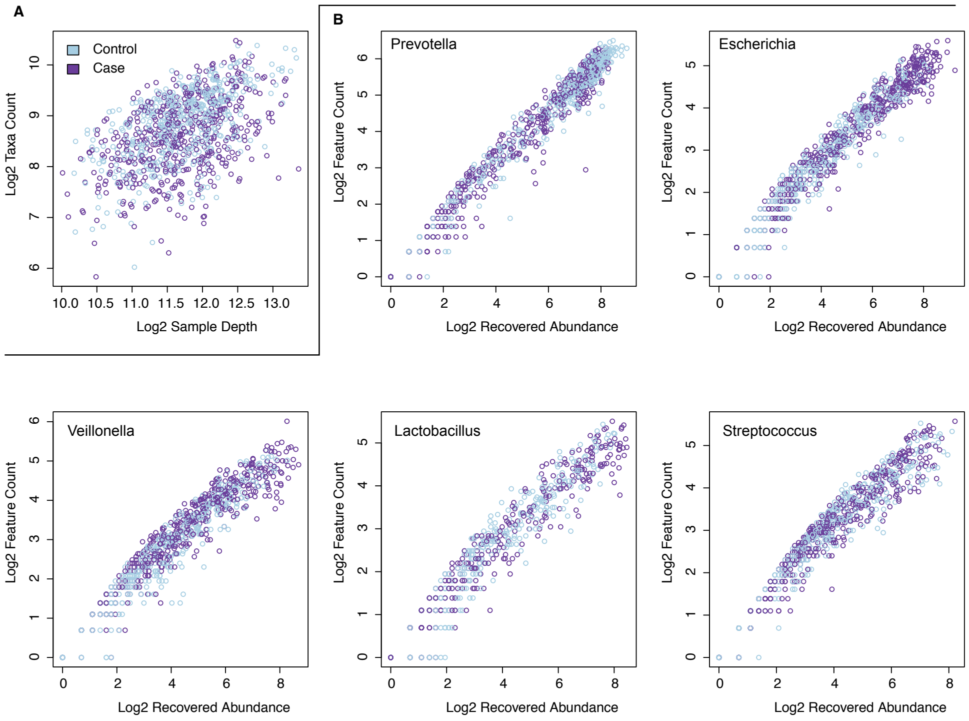


**Figure S1: Within-genus taxa accumulation data are concordant for several genera in 16S surveys.** (A) Taxa accumulations with respect to sample-depth in the diarrheal microbiome dataset. (B) Genus-specific taxa accumulations with respect to the genus recovered abundance for a few genera in the same dataset.

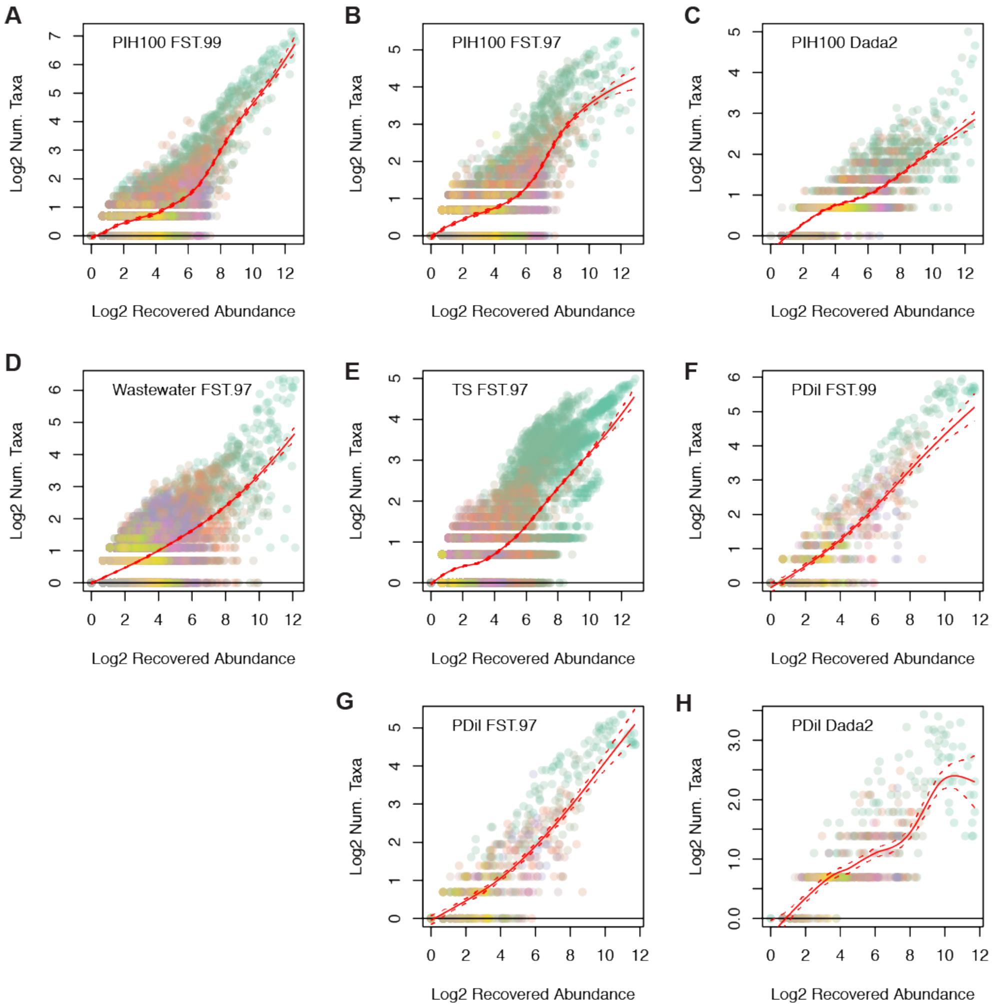


**Figure S2:** Companion to the within-genus Table 1 in main text. Within-genus accumulations are overlaid across genus accumulations in each of the datasets. Smoothing spline $f_R(\cdot)$ within-genus accumulation trends for various datasets. Colors represent distinct genera within each sub-plot.

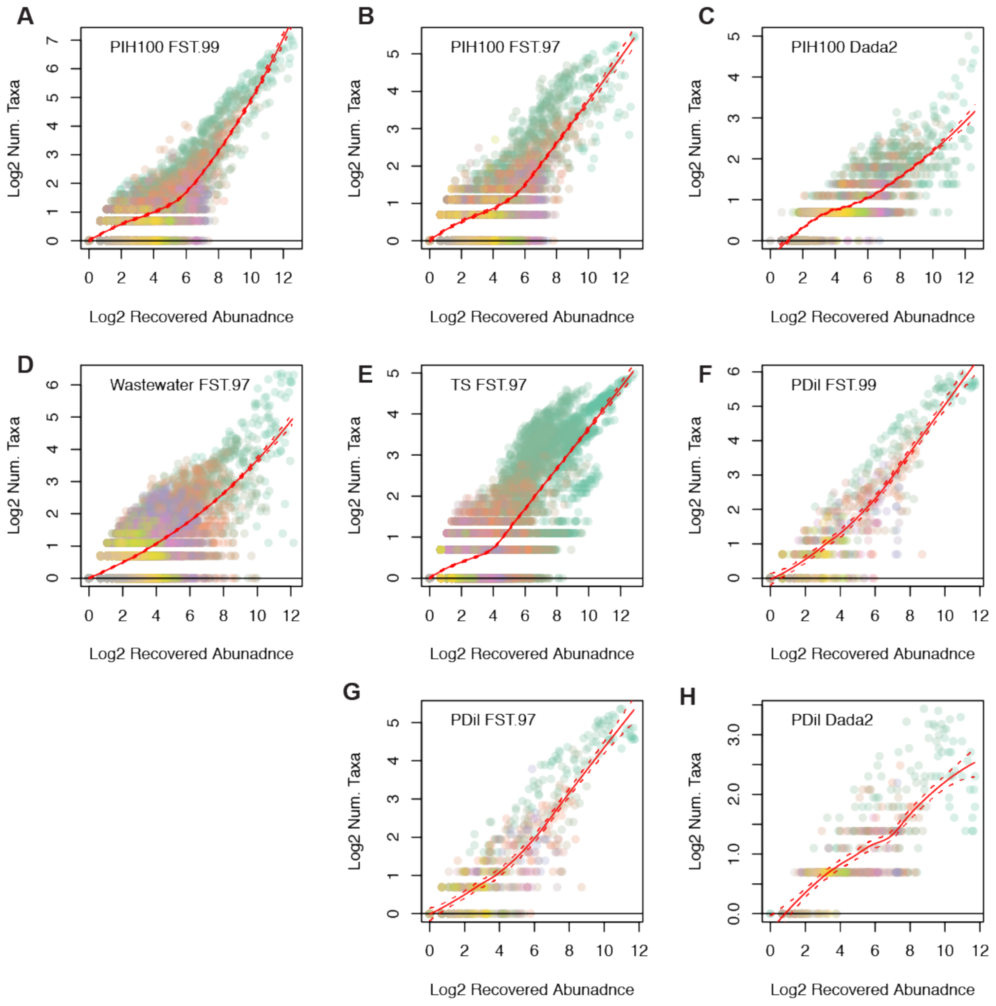


**Figure S3:** Companion to within-genus Table 1 in main text. Within-genus accumulations are overlaid across genus accumulations in each of the datasets. Loess $f_R(\cdot)$ within-genus accumulation trends for various datasets. Colors represent distinct genera within each sub-plot.

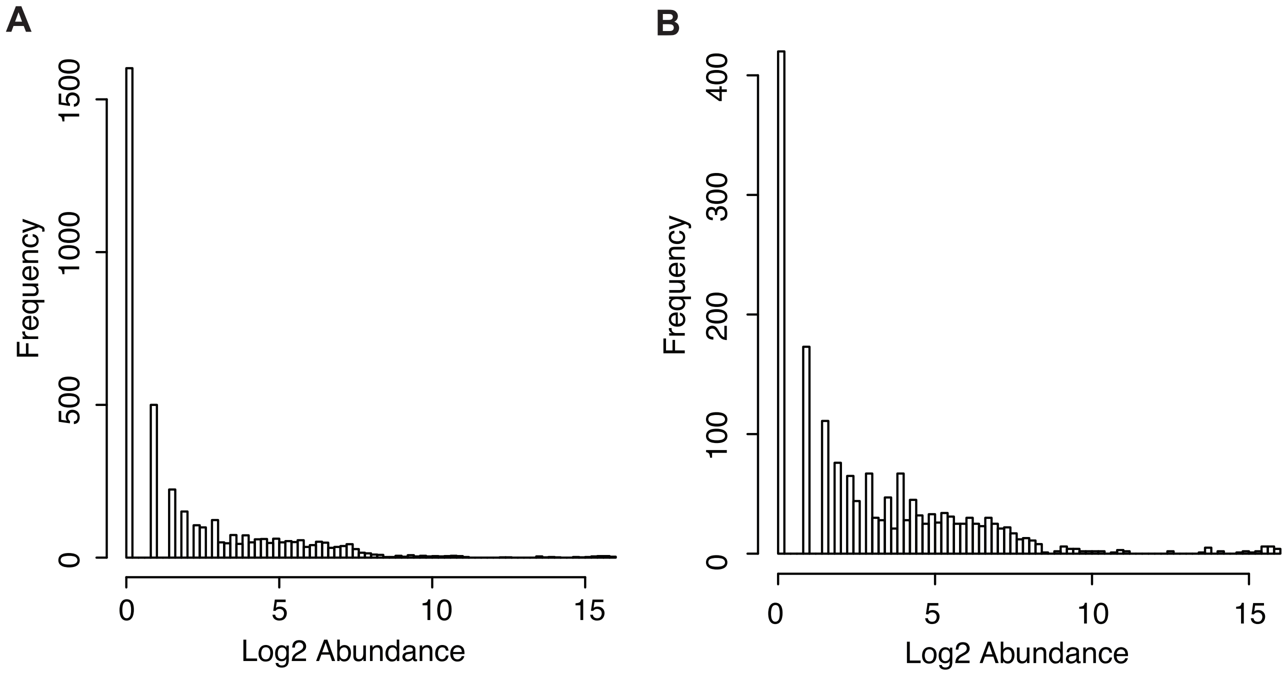


**Figure S4: Non-dominant Pseudomonas taxa discoveries from the dilution study preferentially contribute to the low frequency regime** The Pseudomonas taxon with the largest dataset-wide aggregated count is taken as the dominant input sequence. Frequencies of frequencies of rest of the taxa are plotted for two FST clustering variants, (A) 99% and (B) 97%, including by including all samples from the dataset. The Pseudomonas genera is known to have $\sim 4$ 16S rRNA genes per cell. Up to their sequence conservation, it is possible that larger counts are likely legitimate input sequences.

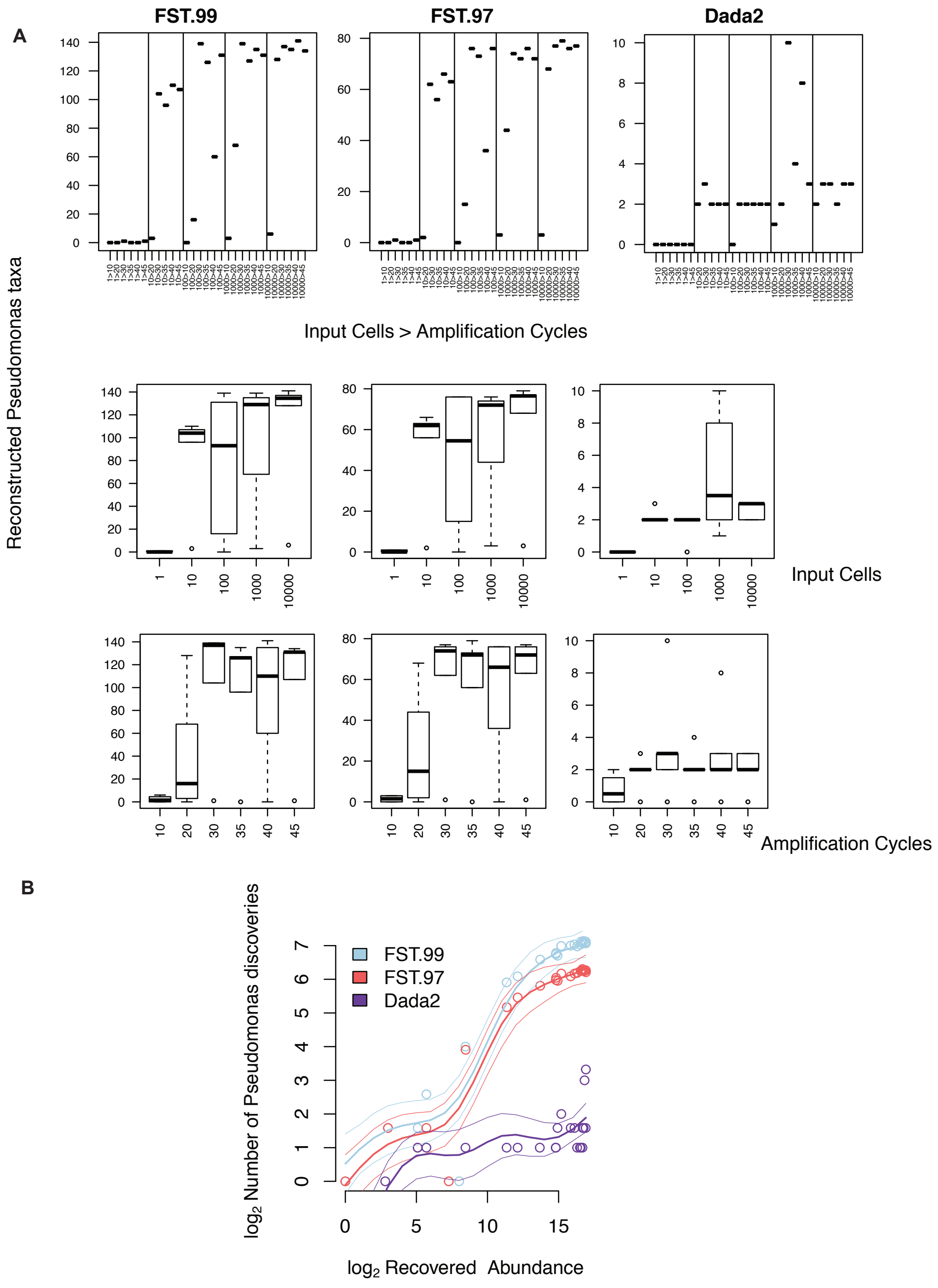


**Figure S5: False microbial discoveries accumulate along the recovered abundance axis** As in main text Fig. 2, but only the taxa that were found to have a significant positive association with experimentally designed input cell numbers, as ascertained by a Poisson regression, are plotted. A caveat is that spurious discoveries have low frequencies, a regime where such tests have low detection power.

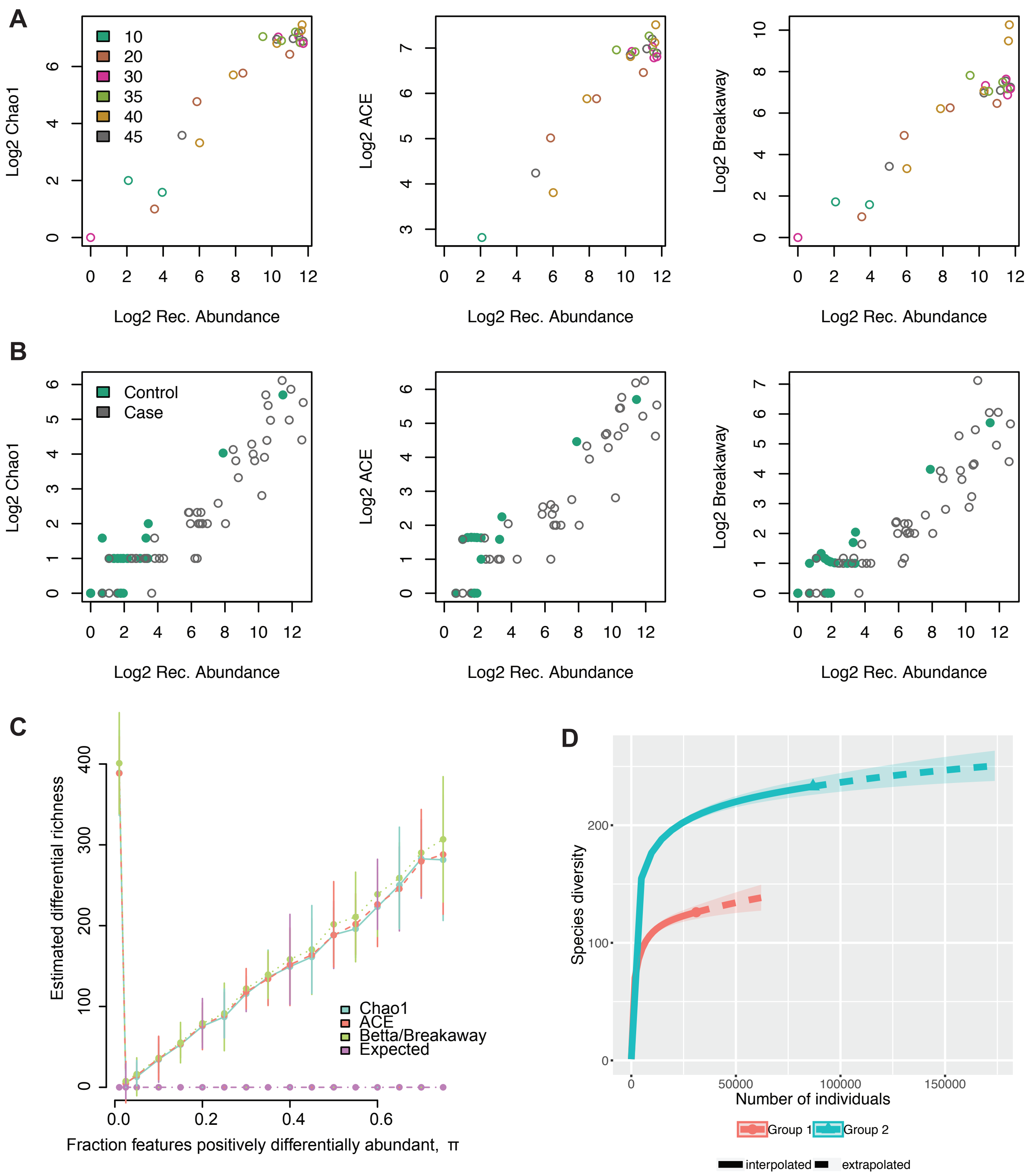


**Figure S6: Systematic bias in some common richness estimators.** The estimated genus-specific richness grows as a function of genus-wide recovered abundance in groups of similar samples. (A) Sample-wise Pseudomonas genus richness estimates from the Pseudomonas experiment. Colors denote amplification cycles. (B) Sample-wise Paenibacillus genus richness estimates from PIH100 dataset. Colors indicate experimental groups. ( C,D) Skewed relative abundance fold change distribution of detected genera across sample-groups lead to differential spurious taxa accumulations, resulting in confounded sample-wide richness inferences. Simulations are performed according to the procedure in supplementary note 3. Asymptotic richness estimators are noted in (C), and interpolated and extrapolated non-asymptotic rarefaction based richness inference for two simulated null samples are shown in (D).

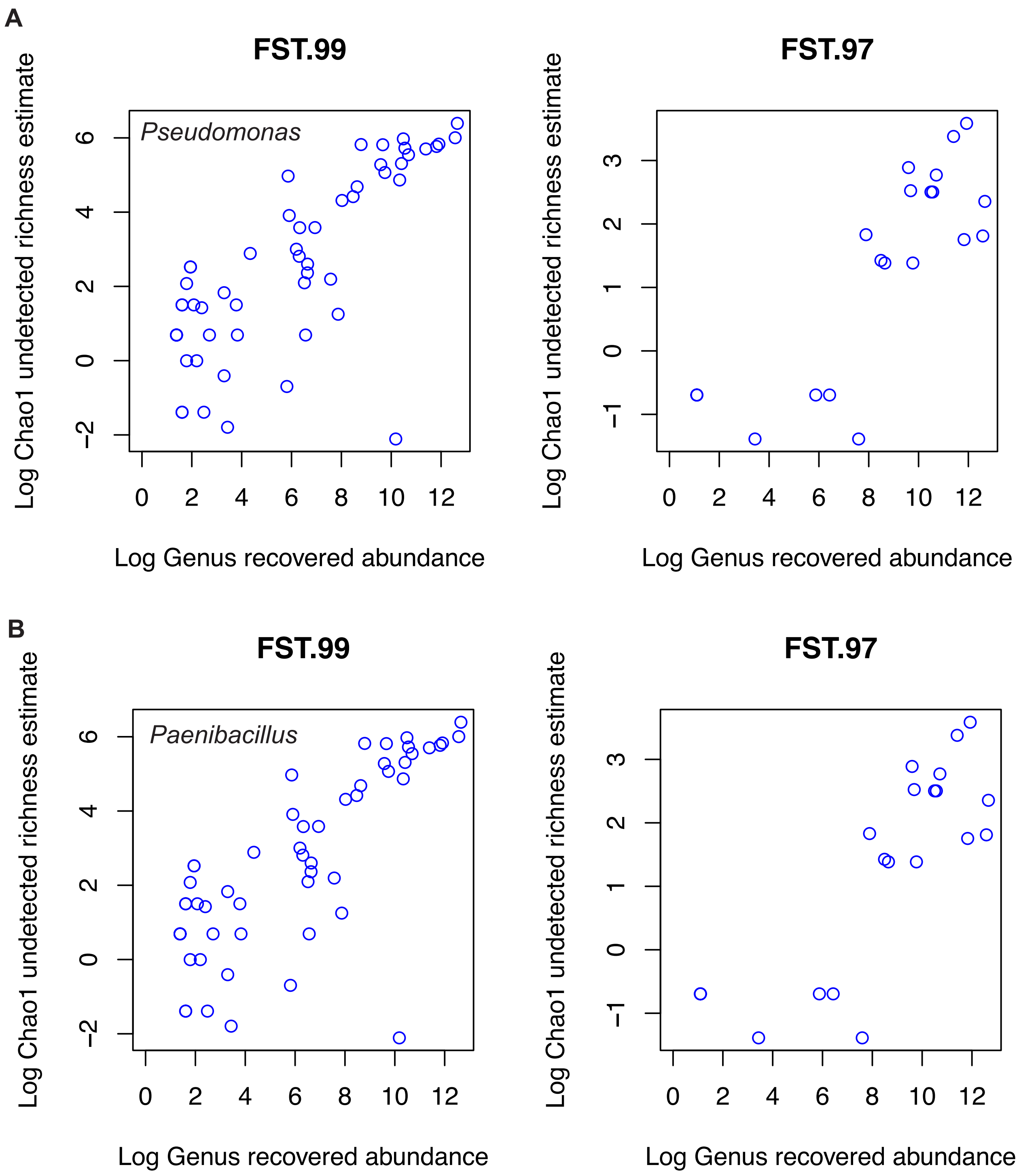


**Figure S7: Chao1 undetected richness estimates grow with genus recovered abundance** The Chao1 undetected richness estimator for the Pseudomonas genus in the Pseudomonas dilution study (top row) and Paenibacillus genus in PIH100 survey (bottom row) for two different settings of sequence similarity thresholds. FST.99 and FST.97 correspond to sequence similarity thresholds of 99% and 97% respectively.

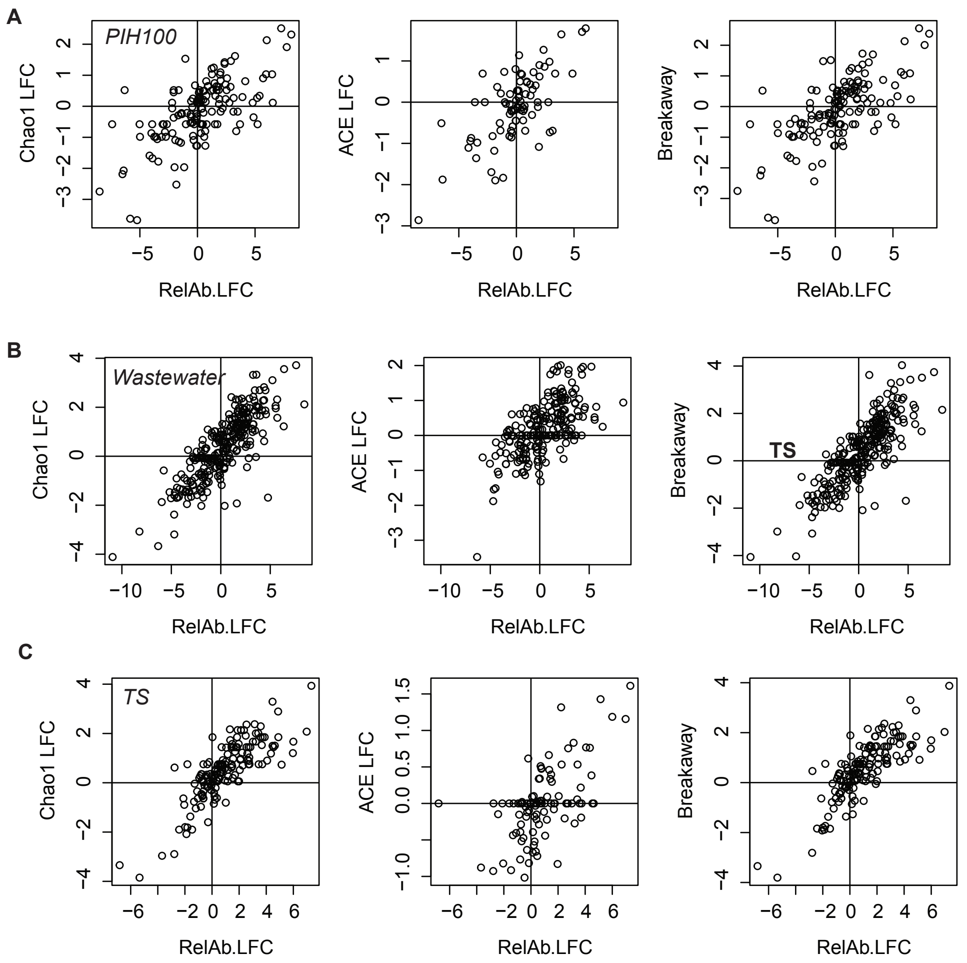


**Figure S8: Differential richness estimators are biased by differential abundance.** Each row corresponds to a two-group comparison for a distinct 16S survey. (A) PIH100 (Case vs. control), (B) Wastewater (Effluent vs. Influent), (C) Long term time series (Post-infection vs. Pre-infection). In each case, the estimated log-fold changes in estimated richness by the noted estimators are plotted against the genus-wise relative abundance log-fold changes across sample groups. In each plot, those genera for which a DR estimate was not available due to estimation problems in respective methods, are not shown.

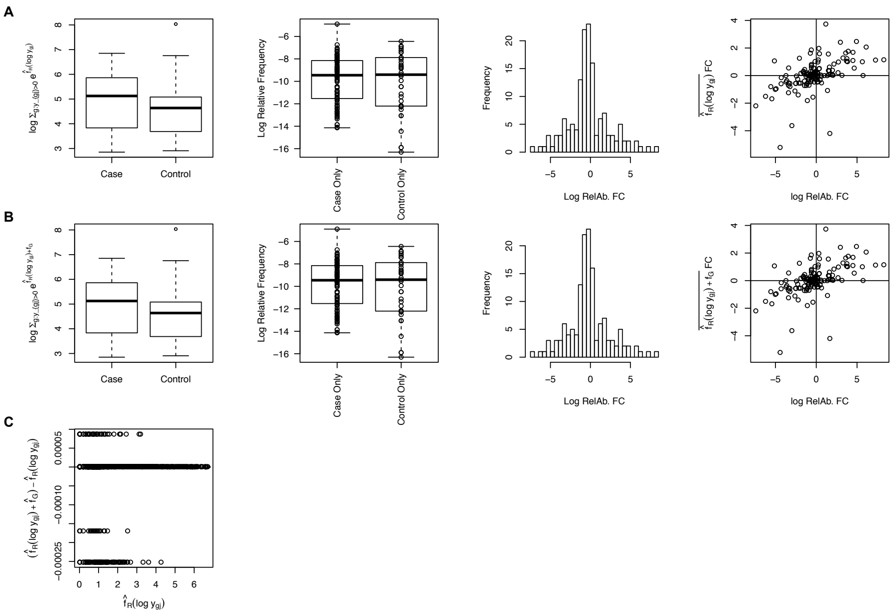

**Figure S9:** A comparison of two technical component estimators for the PIH100 FST99 dataset. (A) without genus contributions, (B) with genus contributions. In both rows, the first column plots the technical predictor in each sample-group. Second column plots the relative abundances of genuses occurring in one of the sample-groups only. The third column plots the histogram of genus-wise case vs. control relative abundance fold changes. The fourth column plots the fold change in the technical predictor against the respective genera's relative abundances. ( C ) plots the two estimators.

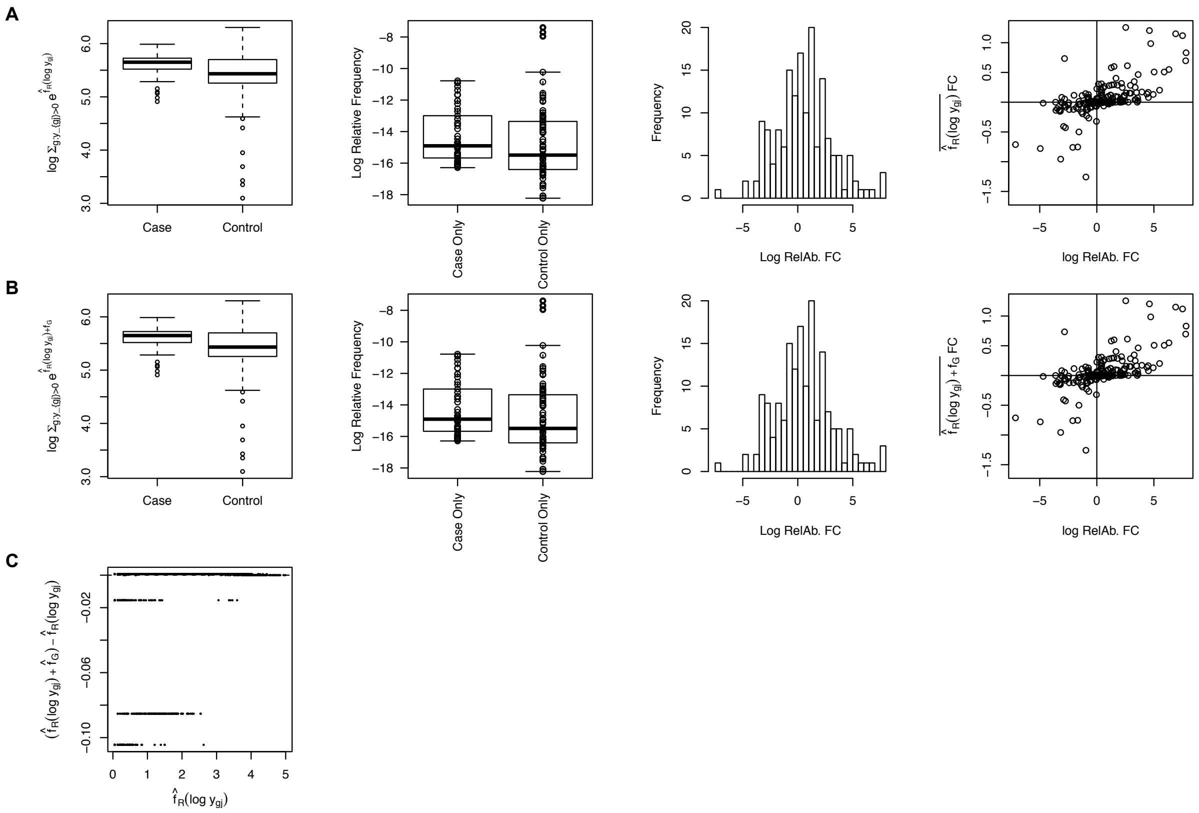


**Figure S10:** A comparison of two technical component estimators for the TS FST97 dataset. Case and control refer to post-infection and pre-infection samples respectively. (A) without genus contributions, (B) with genus contributions. In both rows, the first column plots the technical predictor in each sample-group. Second column plots the relative abundances of genuses occurring in one of the sample-groups only. The third column plots the histogram of genus-wise case vs. control relative abundance fold changes. The fourth column plots the fold change in the technical predictor against the respective genera's relative abundances. ( C ) plots the two estimators.

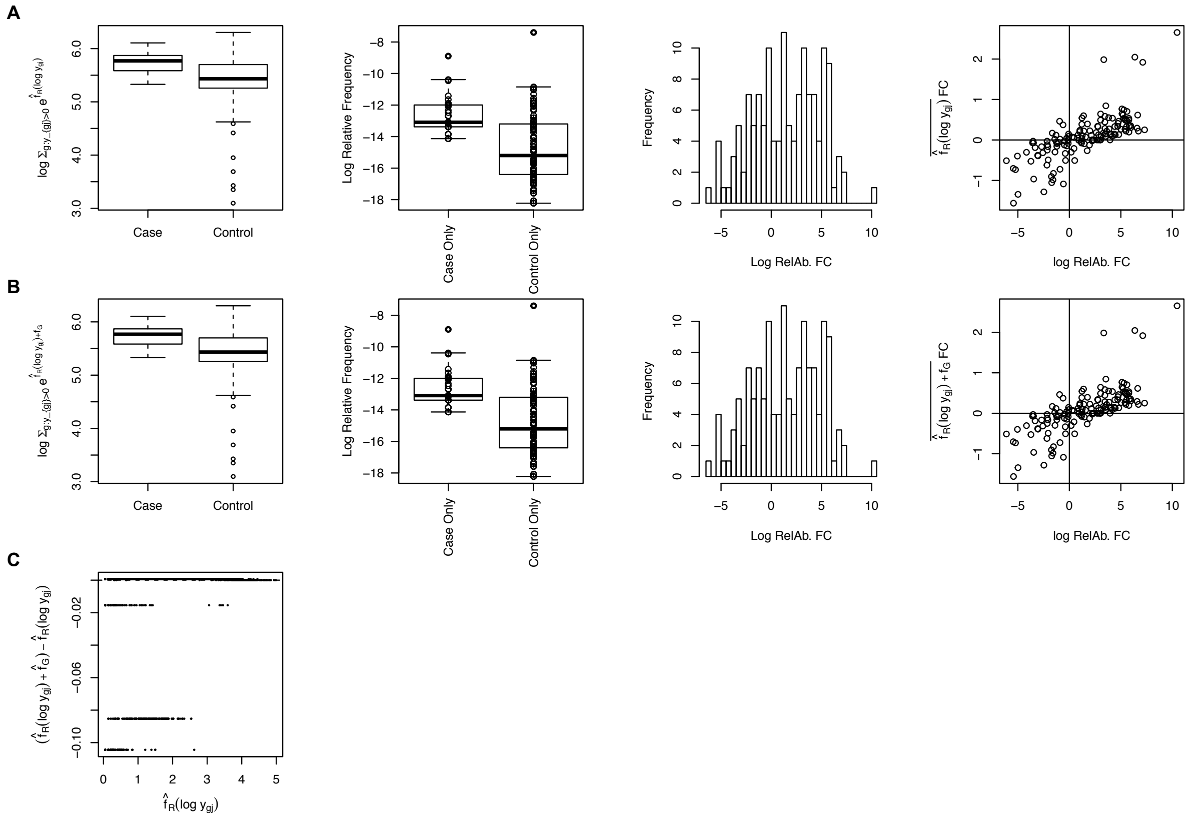


**Figure S11:** A comparison of two technical component estimators for the TS FST97 dataset. Case and control refer to infection and pre-infection samples respectively. (A) without genus contributions, (B) with genus contributions. In both rows, the first column plots the technical predictor in each sample-group. Second column plots the relative abundances of genuses occurring in one of the sample-groups only. The third column plots the histogram of genus-wise case vs. control relative abundance fold changes. The fourth column plots the fold change in the technical predictor against the respective genera's relative abundances. ( C ) plots the two estimators.

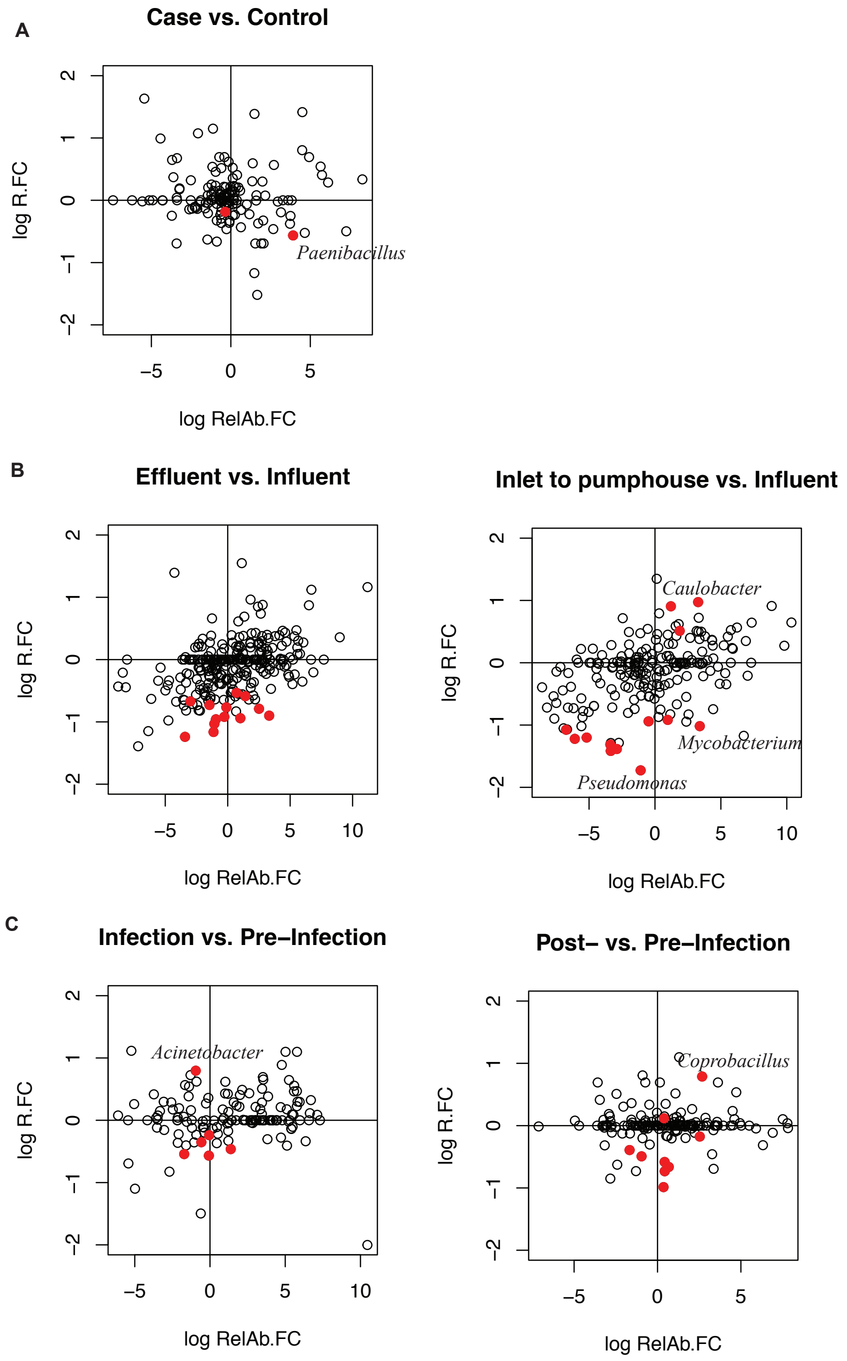


**Figure S12: Genus-specific differential richness inferences from Prokounter.** Each row corresponds to a two-group comparison for a distinct 16S survey. Compare to supplementary Fig. S8. (A) PIH100 (Case vs. control), (B) Wastewater (Effluent vs. Influent), and (Inlet to pumphouse vs Influent), (C) Long term time series (Infection vs. Pre-infection, and Post-infection vs. Pre-infection). In each case, the estimated log-fold changes in estimated richness by the noted estimators are plotted against the genus-wise relative abundance log-fold changes across sample groups. Significant richness differences are colored red.

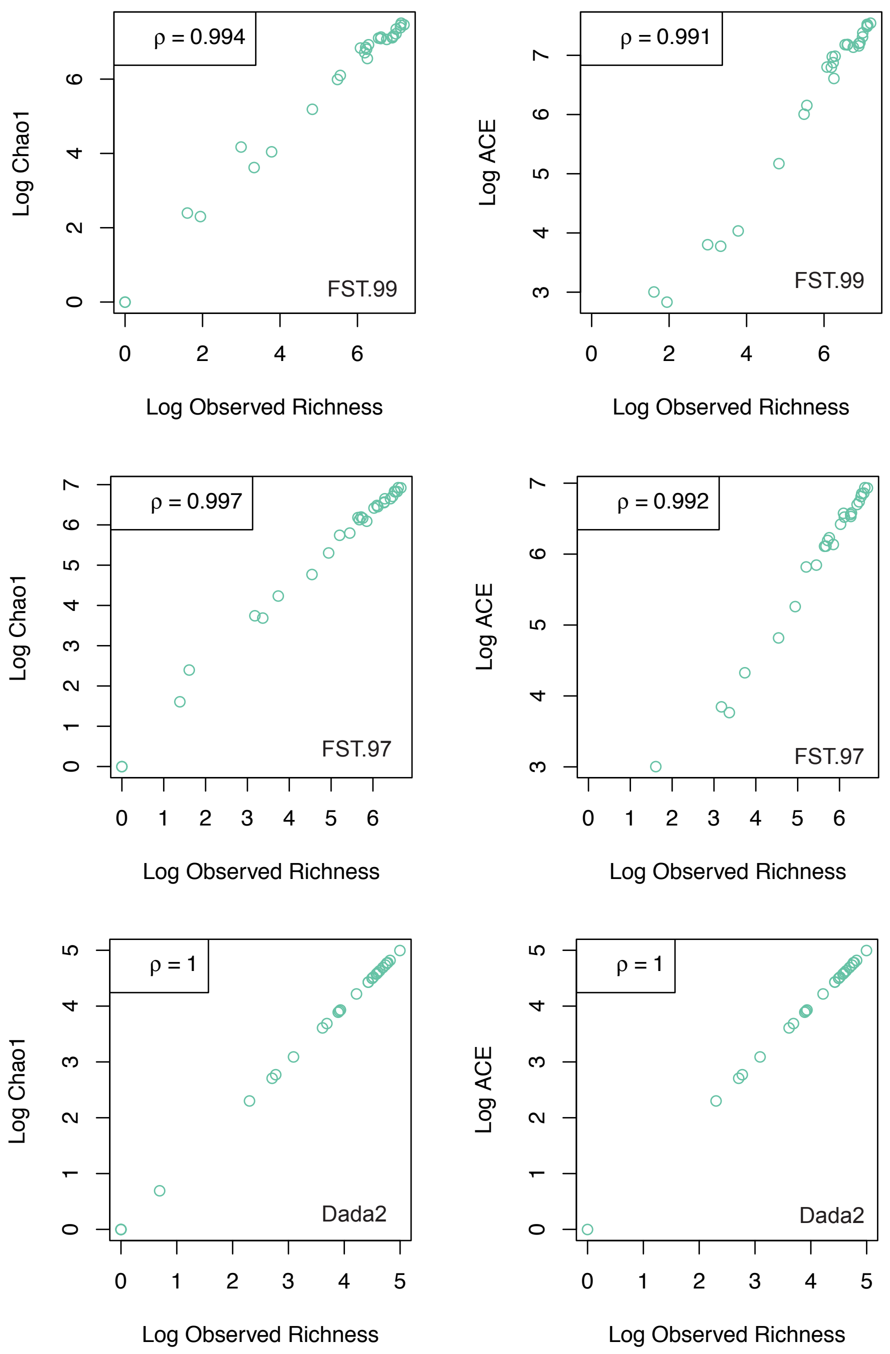


**Figure S13:** Sample-wide richness estimates track observed sample-wide richness in the Pseudomonas dilution study, for various clustering methods mentioned. $\rho$ refers to Pearson correlation.

.

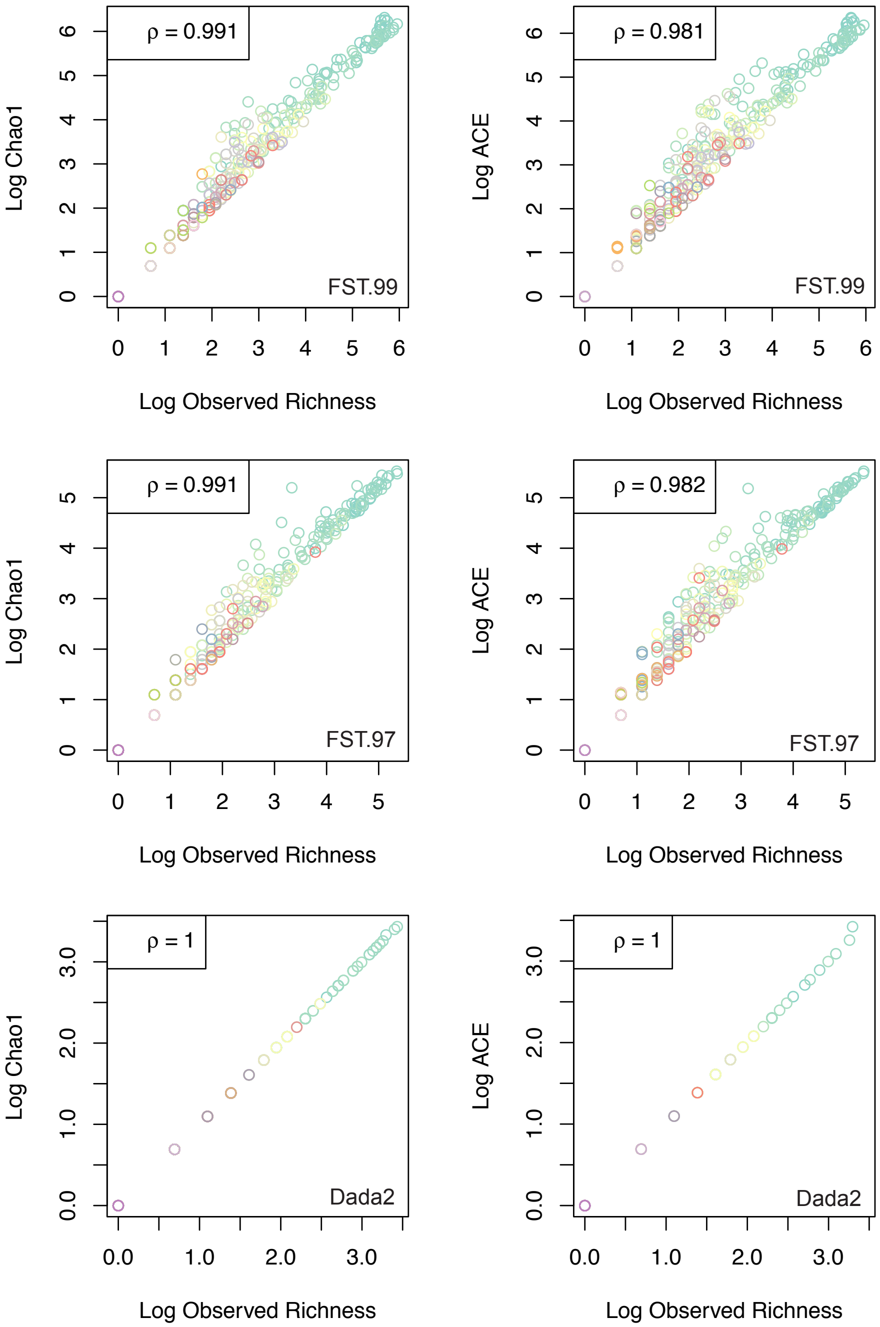


**Figure S14:** Genus-specific richness estimates track genus-specific observed richness values in the Pseudomonas dilution study.

.

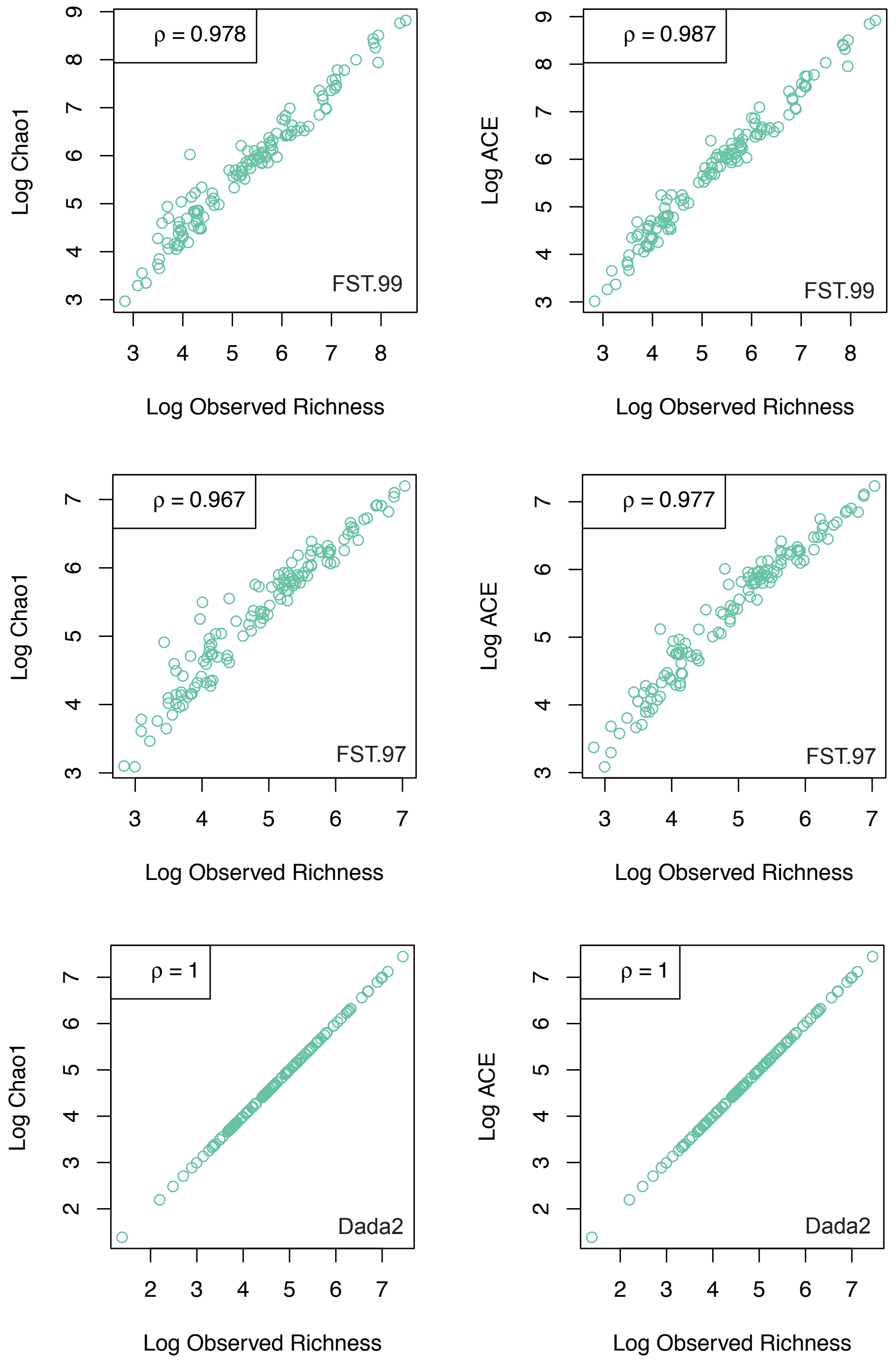


**Figure S15:** Sample-wide richness estimates track observed sample-wide richness in the PIH100 study [1].

.

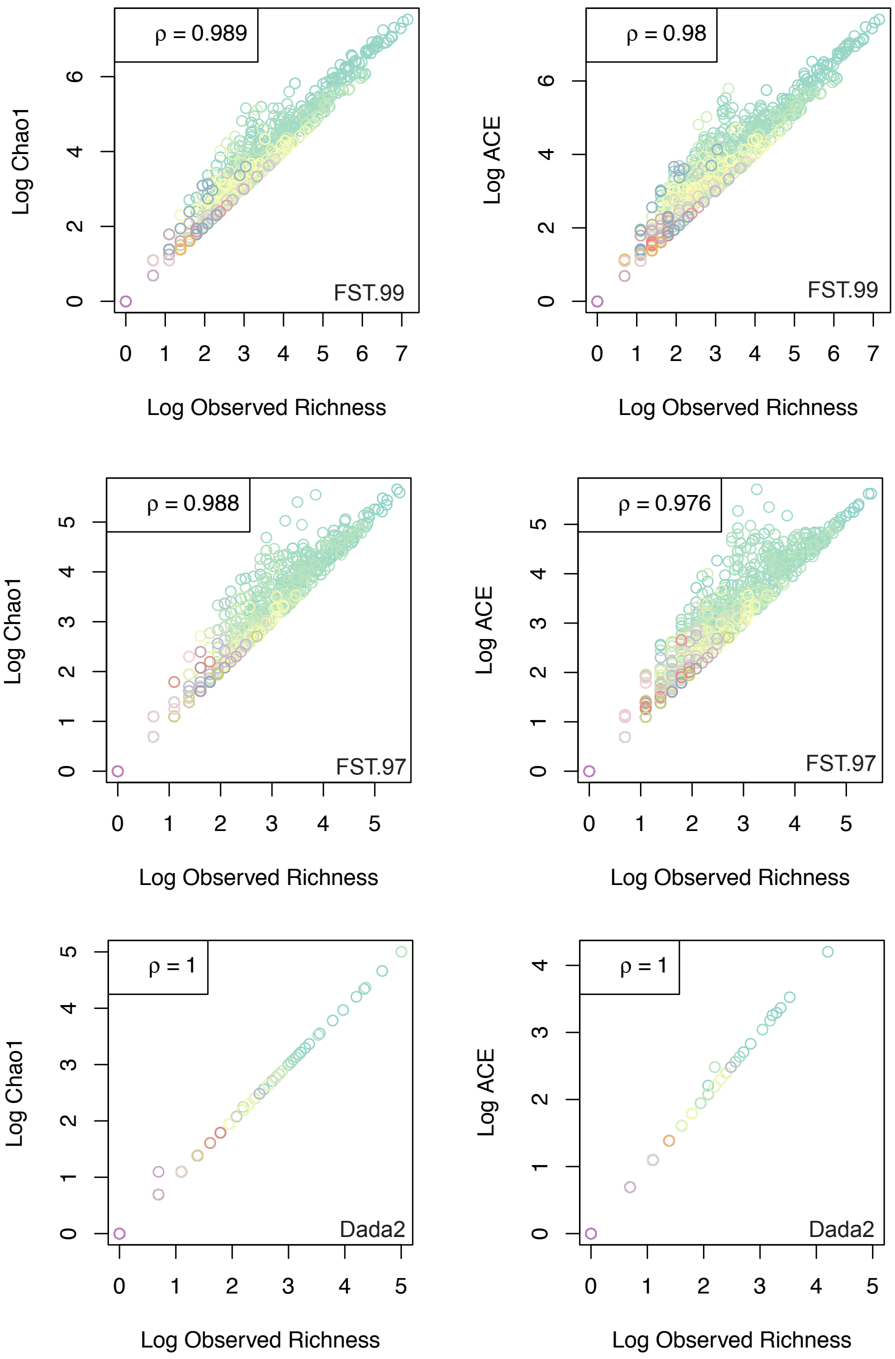


**Figure S16:** Genus-specific richness estimates track genus-specific observed richness values in the PIH100 study [1].

.

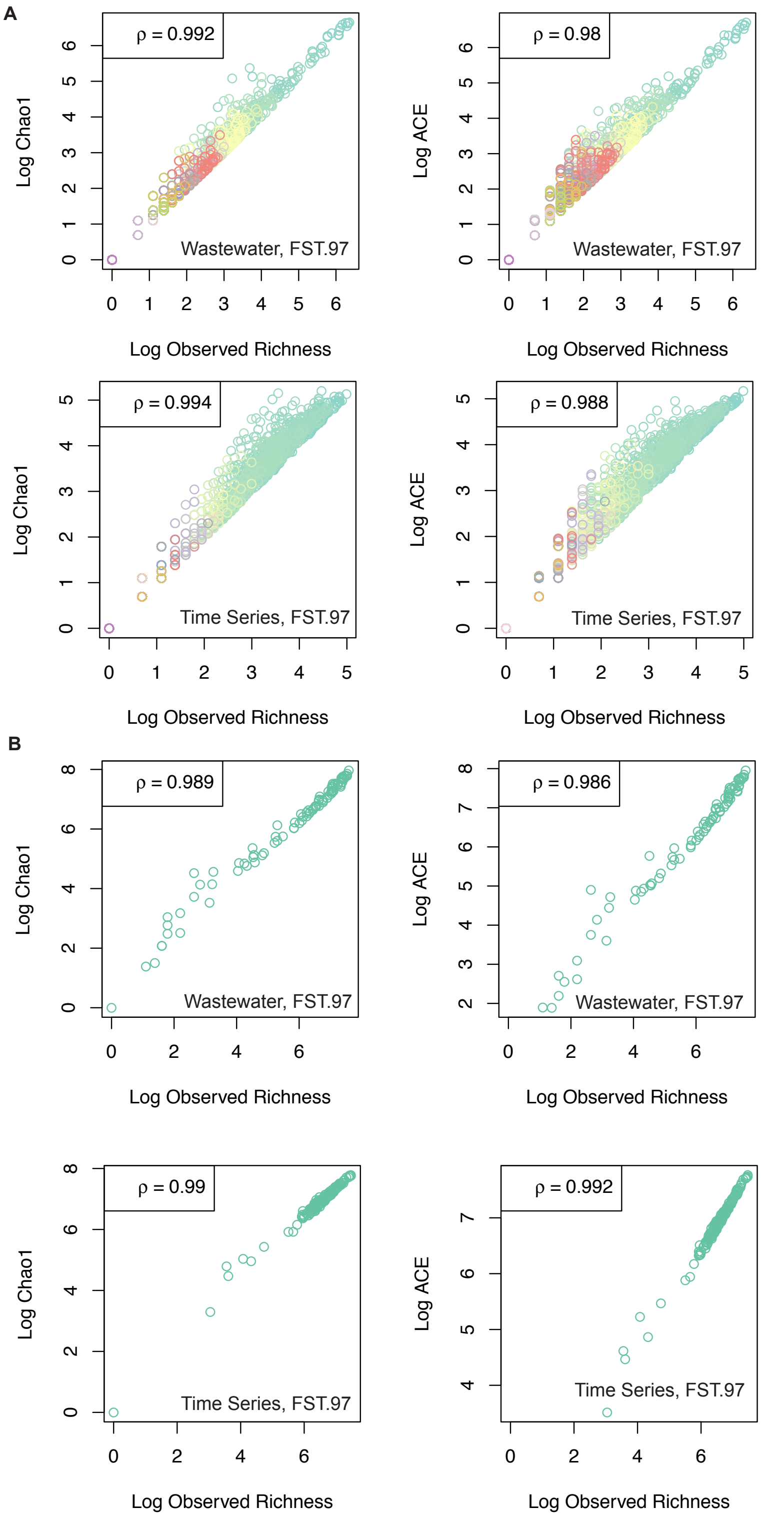


**Figure S17:** (A) Genus-specific richness estimates track genus-specific observed richness values in the wastewater [2] and long-term time series study [3]. (B) Sample-wide richness estimates track observed sample-wide richness in Wastewater and long-term time series study.

.

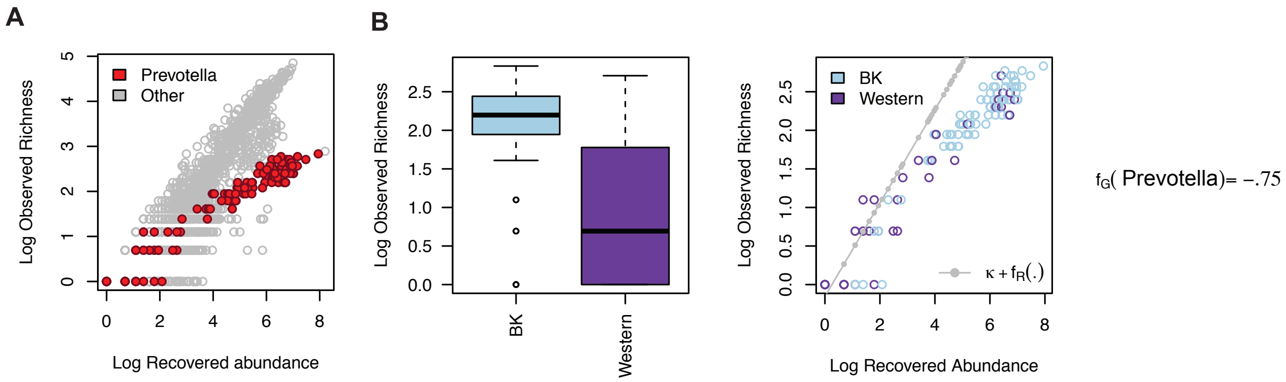


**Figure S18:** (A) Relative to most of the genera, *Prevotella* has a systematically reduced accumulation trend through most of the recovered abundance range in the mouse diet study [4]. (B) This is captured by $f_G$ in the `Prokounter` model. *Prevotella* also has reduced observed richness (number of taxa) in the "Western" group relative to "BK" control diet group. Unless $f_G()$ is incorporated, the abundance dependent technical correction indicated by $f_R()$ is too severe for *Prevotella*, leading one to entirely attribute the observed richness differences between groups to sampling abundance variation.

.

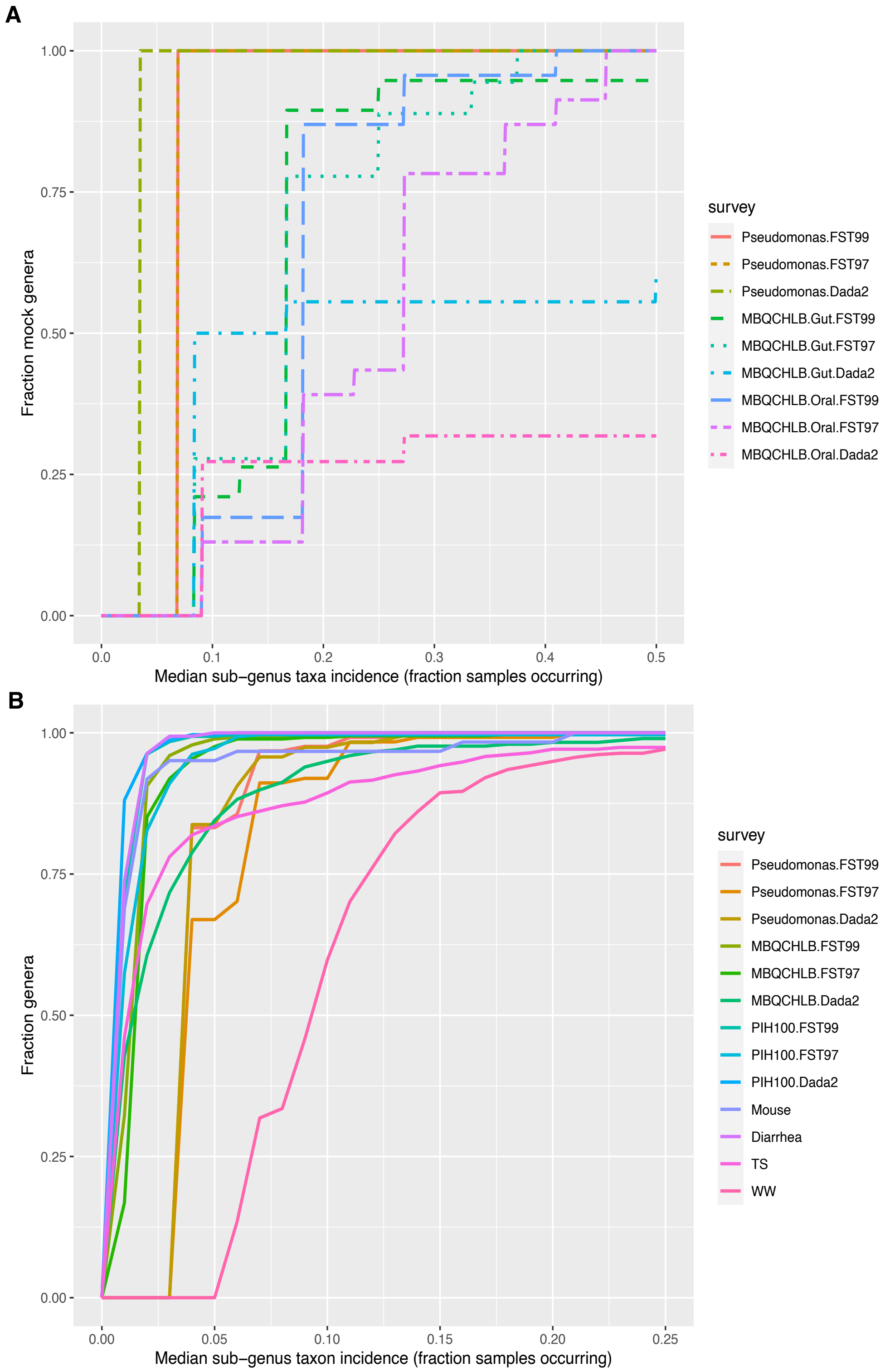


**Figure S19:** Replicability of sub-genus taxa in study samples. For each genus in a given study, its median sub-genus taxa incidence is calculated as the median of the fraction of the study samples in which the genus's taxa occur. This value is plotted along the x-axis. For each incidence value along the x-axis, the y-axis plots the cumulative fraction of genera with at most that medium sub-genus taxa incidence value. (A) restricts this analysis to mock genera in mock datasets only and (B) generalizes this to all represented genera in all datasets considered in this paper.

.

### Supplementary Tables

### Supplementary Tables

| Dataset | Year | 16S segment, Sequencing& Clustering | Number of detected genera | Optimal number of clusters |
|---|---|---|---|---|
| Mouse [1, 2] | 2009 | V2, 454, FST.97 | 60 | 2 |
| Diarrhea [3] | 2014 | V12, 454, FST.99 | 163 | 2 |
| Time series [4] | 2014 | V4, GAIIx, FST.97 | 309 | 3 |
| Wastewater [5] | 2018 | V34, MiSeq, FST.99 | 412 | 4 |
| MBQC-HLB$^{(97)}$ [6] | 2017 | V4, MiSeq, FST.97 | 370 | 3 |
| MBQC-HLB$^{(99)}$ [6] | 2017 | V4, MiSeq, FST.99 | 373 | 3 |
| MBQC-HLB$^{(D)}$ [6] | 2017 | V4, MiSeq, Dada2 | 298 | 2 |
| PIH100$^{(97)}$ [7] | 2020 | V12, MiSeq, FST.97 | 288 | 6 |
| PIH100$^{(99)}$ [7] | 2020 | V12, MiSeq, FST.97 | 318 | 5 |
| PIH100$^{(D)}$ [7] | 2020 | V12, MiSeq, Dada2 | 288 | 6 |
| Pseudomonas$^{(97)}$ | 2021 | V12, MiSeq, FST.97 | 124 | 2 |
| Pseudomonas$^{(99)}$ | 2021 | V12, MiSeq, FST.99 | 125 | 8 |
| Pseudomonas$^{(D)}$ | 2021 | V12, MiSeq, Dada2 | 117 | 5 |

**Table S1: Within-genus taxa accumulation data organize into very few accumulation patterns.** For each survey, its within-genus taxa accumulation data is organized into two columns as follows. The first column holds, the raw recovered abundance values for each genus for all samples, and the second column holds the corresponding number of observed taxa. The recovered abundance values were binned into 25 windows, with comparable number of observations (i.e., rows in the two column table) in each window. A data matrix was next generated with rows representing genus, columns representing each of the 25 windows, and each cell corresponding to the mean number of taxa observed within the cell. A hierarchical clustering tree was constructed on this set of accumulation patterns, and the optimal number of clusters corresponding to the minimal silhouette width was obtained (column 5) out of the possible number of clusters (number of genera, column 4). These numbers are in good agreement with within-genus taxa accumulation data visualizations.

| Dataset | Year | 16S segment, Sequencing& Clustering | Pseudo $R^2_{\log\ depth}$ | Pseudo $R^2_{\log\ depth+design}$ | $AIC_{\log\ depth}$ | $AIC_{\log\ depth+design}$ |
|---|---|---|---|---|---|---|
| Mouse [1, 2] | 2009 | V2, 454, FST.97 | 99.04% | 99.14% | $1.3853 \times 10^3$ | $1.3230 \times 10^3$ |
| Diarrhea [3] | 2014 | V12, 454, FST.99 | 99.92% | 99.93% | $1.2727 \times 10^4$ | $1.2578 \times 10^4$ |
| Time series [4] | 2014 | V4, GAIIx, FST.97 | 99.95% | 99.95% | $2.1964 \times 10^3$ | $2.1744 \times 10^3$ |
| Wastewater [5] | 2018 | V34, MiSeq, FST.99 | 99.94% | 99.95% | $7.2713 \times 10^2$ | $7.1527 \times 10^2$ |
| MBQC-HLB$^{(97)}$ [6] | 2017 | V4, MiSeq, FST.97 | 99.83% | 99.85% | $2.7512 \times 10^3$ | $2.6998 \times 10^3$ |
| MBQC-HLB$^{(99)}$ [6] | 2017 | V4, MiSeq, FST.99 | 99.98% | 99.99% | $2.9250 \times 10^3$ | $2.8966 \times 10^3$ |
| MBQC-HLB$^{(D)}$ [6] | 2017 | V4, MiSeq, Dada2 | 99.67% | 99.77% | $2.0710 \times 10^3$ | $1.9483 \times 10^3$ |
| PIH100$^{(97)}$ [7] | 2020 | V12, MiSeq, FST.97 | 99.83% | 99.87% | $1.3833 \times 10^3$ | $1.3133 \times 10^3$ |
| PIH100$^{(99)}$ [7] | 2020 | V12, MiSeq, FST.99 | 99.95% | 99.95% | $1.4208 \times 10^3$ | $1.3853 \times 10^3$ |
| PIH100$^{(D)}$ [7] | 2020 | V12, MiSeq, Dada2 | 99.46% | 99.58% | $1.1647 \times 10^3$ | $1.1074 \times 10^3$ |
| Pseudomonas$^{(97)}$ | 2021 | V12, MiSeq, FST.97 | 99.92% | 99.95% | $3.2197 \times 10^2$ | $2.900 \times 10^2$ |
| Pseudomonas$^{(99)}$ | 2021 | V12, MiSeq, FST.99 | 99.61% | 99.98% | $3.4318 \times 10^2$ | $3.1313 \times 10^2$ |
| Pseudomonas$^{(D)}$ | 2021 | V12, MiSeq, Dada2 | 99.60% | 99.69% | 244.77 | 234.77 |

**Table S2: Relative to study variables, sample depth explains bulk of the systematic variation in a 16S survey's *sample-wide* taxa accumulations.** For each 16S survey dataset mentioned in column 1, the year of publication is listed in column 2, the partial 16S segment targeted, machine technology and sequence clustering approach used are specified in column 3. McFadden's pseudo-$R^2$ for explaining genus-specific taxa accumulations with two negative binomial regressions (NB) are listed in columns 4 and 5. The fourth column is obtained when the NB regression includes logged sampled depth alone as predictor. The fifth column additionally includes the experimental design matrix for each dataset as predictors.

| Dataset | Year | 16S segment, Sequencing& Clustering | Pseudo $R^2_{\log\ depth}$ | Pseudo $R^2_{\log\ depth+design}$ | $AIC_{\log\ depth}$ | $AIC_{\log\ depth}$ |
|---|---|---|---|---|---|---|
| Mouse [1, 2] | 2009 | V2, 454, FST.97 | 83.62% | 96.69% | $1.7443 \times 10^4$ | $1.2576 \times 10^4$ |
| Diarrhea [3] | 2014 | V12, 454, FST.99 | 87.45% | 94.66% | $1.8648 \times 10^5$ | $1.5884 \times 10^5$ |
| Time series [4] | 2014 | V4, GAIIx, FST.97 | 77.52% | 97.22% | $5.2421 \times 10^4$ | $3.4657 \times 10^4$ |
| Wastewater [5] | 2018 | V34, MiSeq, FST.99 | 69.45% | 90.42% | $3.4313 \times 10^4$ | $2.7412 \times 10^4$ |
| MBQC-HLB$^{(97)}$ [6] | 2017 | V4, MiSeq, FST.97 | 84.28% | 92.65% | $1.5078 \times 10^5$ | $1.3051 \times 10^5$ |
| MBQC-HLB$^{(99)}$ [6] | 2017 | V4, MiSeq, FST.99 | 89.15% | 94.33% | $1.6308 \times 10^5$ | $1.4528 \times 10^5$ |
| MBQC-HLB$^{(D)}$ [6] | 2017 | V4, MiSeq, Dada2 | 40.72% | 73.48% | $4.2232 \times 10^4$ | $3.5720 \times 10^4$ |
| PIH100$^{(97)}$ [7] | 2020 | V12, MiSeq, FST.97 | 80.67% | 91.23% | $1.3833 \times 10^3$ | $1.3133 \times 10^3$ |
| PIH100$^{(99))}$ [7] | 2020 | V12, MiSeq, FST.99 | 89.86% | 95.10% | $1.4210 \times 10^3$ | $1.3853 \times 10^3$ |
| PIH100$^{(D)}$ [7] | 2020 | V12, MiSeq, Dada2 | 74.48% | 82.53% | $1.1647 \times 10^3$ | $1.1073 \times 10^3$ |
| Pseudomonas$^{(97)}$ | 2021 | V12, MiSeq, FST.97 | 93.34% | 98.54% | $4.081 \times 10^3$ | $3.2653 \times 10^3$ |
| Pseudomonas$^{(99)}$ | 2021 | V12, MiSeq, FST.99 | 89.13% | 97.70% | $4.5456 \times 10^3$ | $3.7133 \times 10^3$ |
| Pseudomonas$^{(D)}$ | 2021 | V12, MiSeq, Dada2 | 75.98% | 91.25% | $2.2810 \times 10^3$ | $2.0138 \times 10^3$ |

**Table S3: Relative to study variables, sample-depth explains bulk of the systematic variation in a 16S survey's *genus-specific* taxa accumulations.** For each 16S survey dataset mentioned in column 1, the year of publication is listed in column 2, the partial 16S segment targeted, machine technology and sequence clustering approach used are specified in column 3. McFadden's pseudo-$R^2$ for explaining sample-wide taxa accumulations with two negative binomial regressions (NB) are listed in columns 4 and 5. The fourth column is obtained when the NB regression includes logged genus recovered abundances alone as predictor. The fifth column additionally includes the experimental design matrix for each dataset as predictors. Six and seventh columns indicate the respective Akaike Information Criteria.
